# A mechanosensitive RhoA pathway that protects epithelia against acute tensile stress

**DOI:** 10.1101/281154

**Authors:** Bipul R Acharya, Alexander Nestor-Bergmann, Xuan Liang, Srikanth Budnar, Oliver E. Jensen, Zev Bryant, Alpha S. Yap

**Affiliations:** Division of Cell Biology and Molecular Medicine, Institute for Molecular Bioscience, The University of Queensland, St. Lucia, Brisbane Australia 4072; Department of Physiology, Development and Neuroscience, University of Cambridge, Cambridge CB2 3DY, UK; School of Mathematics, University of Manchester, Manchester M13 9PL, UK.; Department of Bioengineering, Stanford University and Department of Structural Biology, Stanford University School of Medicine, Stanford, CA 94305, USA

**Keywords:** Epithelia, mechanotransduction, tension, RhoA, Myosin VI

## Abstract

Adherens junctions are tensile structures that couple epithelial cells together. Junctional tension can arise from cell-intrinsic application of contractility or from the cell-extrinsic forces of tissue movement. In all these circumstances, it is essential that epithelial integrity be preserved despite the application of tensile stress. In this study, we identify junctional RhoA as a mechanosensitive signaling pathway that responds to epithelial stress. The junctional specificity of this response is mediated by the heterotrimeric protein Gα12, which is recruited by E-cadherin and, in turn, recruits p114 RhoGEF to activate RhoA. Further, we identify Myosin VI as a key mechanosensor, based on its intrinsic capacity to anchor E-cadherin to F-actin when exposed to tensile load. Tension-activated RhoA signaling was necessary to preserve epithelial integrity, which otherwise undergoes fracture when monolayer stress is acutely increased by calyculin. Paradoxically, this homeostatic RhoA signaling pathway increases junctional actomyosin, a contractile response that might be expected to itself promote fracture. Simulations of a vertex-based model revealed that the protective effect of RhoA signaling can be explained through increased yield limit at multicellular vertices, where experiments showed p114 RhoGEF was necessary to increase E-cadherin and promote actin assembly and organization.

## Introduction

Epithelia are the fundamental tissue barriers of the body. Epithelial integrity requires that cells be coupled together by specialized cell-cell junctions that can resist mechanical stresses applied to tissues. Of these junctions, a notable role is played by cadherin-based adherens junctions (AJ), which bear mechanical tension and preserve tissue integrity (Harris, et al., 2014; Levine, et al., 1994). This tension arises both from intrinsic forces, where the contractile actomyosin cytoskeleton is coupled to cadherin adhesion; and extrinsic forces such as those associated with animal locomotion and gut peristalsis (Charras and Yap, 2018). Intrinsic junctional tension is evident even in apparently quiescent monolayers (Ratheesh, et al., 2012); but tension is also stimulated to mediate a wide range of morphogenetic processes, from cell-cell rearrangements (Lecuit and Lenne, 2007) to cell extrusion (Michael, et al., 2016). Thus, increases in junctional tension often serve fundamental physiological roles. However, in all these circumstances it is necessary for cell-cell cohesion to be preserved despite the application of force.

One potential solution to this challenge is for cells to possess mechanisms that can sense junctional stress and elicit proportionate homeostatic responses that preserve tissue integrity. Indeed, it is increasingly evident that mechanosensitive molecular mechanisms exist at AJ, and a wide range of cell signals are found there that can modulate adhesion and the cytoskeleton, key elements for junctional integrity (Yap, et al., 2017; Pruitt, et al., 2014). For example, catch bonds mediate the interaction between the cadherin molecular complex and actin filaments (Buckley, et al., 2014), while Rho family GTPases and Src family kinases are found to signal at junctions (Gomez, et al., 2015; Ratheesh, et al., 2012). Despite this growing wealth of candidates, we have yet to holistically characterize a pathway that might maintain junctional integrity against stress. For this, we would need to identify key mechanosensitive receptors; the signaling pathways that are elicited; and the effector mechanisms that preserve epithelial integrity.

We now report such a pathway that is necessary to preserve epithelial integrity against both intrinsic and extrinsic stress. We show that junctional RhoA signaling is activated in response to mechanical stress. We identify the mechanosensitive Myosin VI as a key force-sensor that stabilizes E-cadherin to permit recruitment of the RhoA activators, Gα12 and p114Rho GEF. Surprisingly, despite increasing junctional actomyosin, this pathway prevents epithelial junctions from fracturing upon application of stress. Combining a predictive mechanical model for epithelial junctions with experiments, we find that this can be explained by an increase in yield limit due to RhoA-dependent stabilization of E-cadherin at multicellular vertices.

## Results

### Mechanical stress activates RhoA at AJ

We first combined computational prediction and experiment to understand how increasing contractility might affect monolayers. Intrinsic contractile stress was increased by treating confluent Caco-2 monolayers with calyculin A (henceforth calyculin; 20 nM, 12 min; Figure 1A) which stimulates non-muscle myosin (NMII) by inhibiting the PP2A subunit of myosin phosphatase. Western analysis confirmed that ppMLC levels were increased (Figures S1A and S1B) and immunofluorescence microscopy showed that ppMLC was increased both at junctions and in the cytoplasm (Figures 1B and 1C).

**Figure 1.**
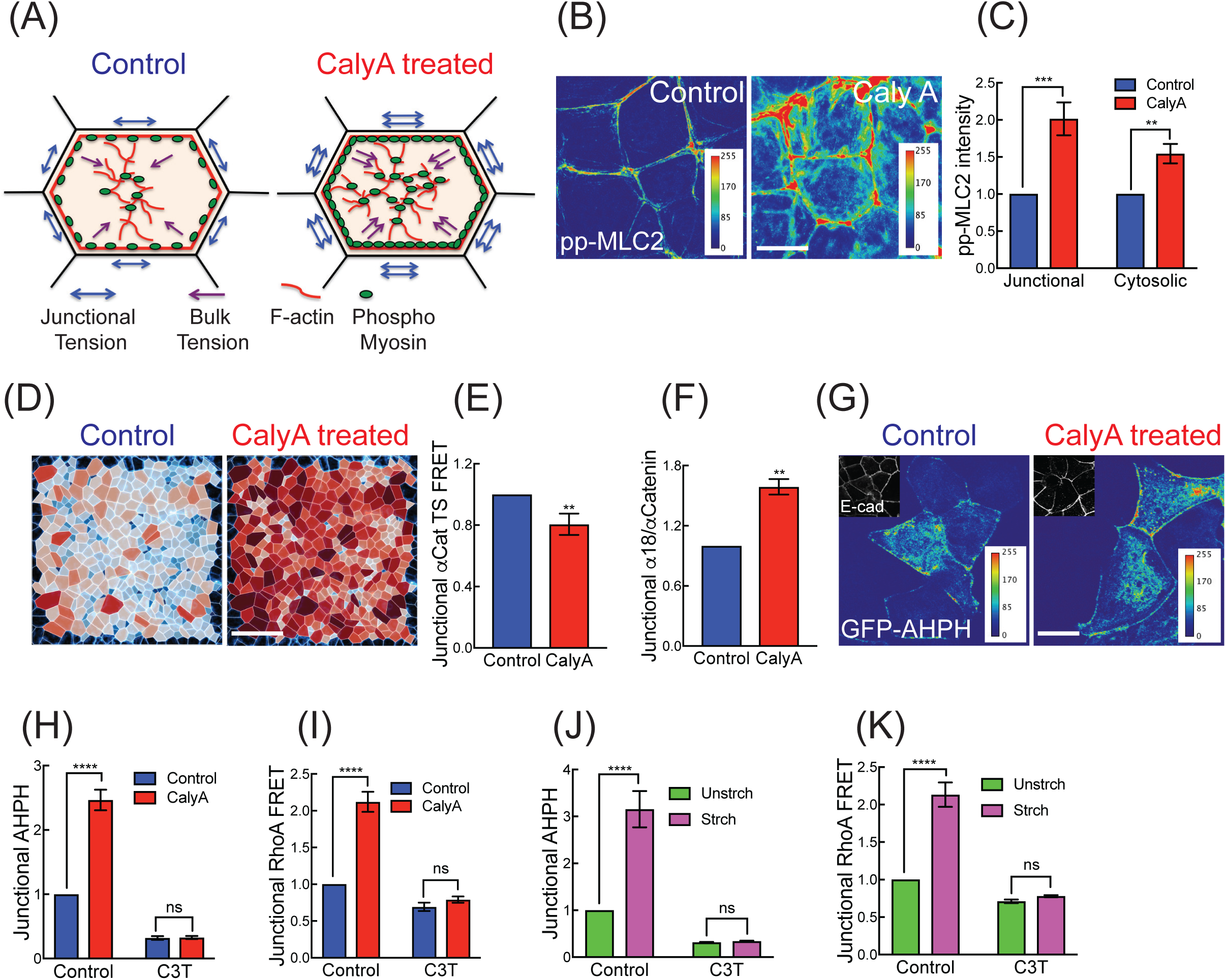
Tensile monolayer stress activates RhoA at the zonula adherens. (A) Cartoon of contractile activation by calyculin A. (B, C) Calyculin A increases active Myosin II (pp-MLC2, pT18/S19) at junctions and in the cytosol: representative images (B) and quantitation (C). (D) Segmented cells on E-cadherin GFP monolayer showing predicted distribution of effective cell pressures, *P*^eff^, from an experiment before (LEFT; *ϕ*_A_ = 1, *ϕ*_P_ = 1) and after (RIGHT; *ϕ*_A_ = 0.75, *ϕ*_P_= 1.25) treatment with Calyculin. Cells in darker red (blue) are predicted to be under higher net tension (compression). *ϕ*_A_and *ϕ*_P_ represent fractional change in preferred area and cortical stiffness respectively and are defined in the Computational Supplement. (E, F) Calyculin A increases molecular-level tension across αE-Catenin, measured with (E) a FRET-based αE-Cat tension sensor (normalized to untreated control) and (F) immunofluorescence for α18 mAb (normalized to total junctional αE-Catenin). (G, H) Calyculin A increases junctional AHPH: representative images for GFP-AHPH and E-cadherin (G) and quantitation (normalized to cytosolic AHPH, H). C3-transferase (C3T) was used as a negative control. (I) Effect of calyculin A on junctional RhoA-FRET index (normalized to untreated control). (J, K) Effect of mechanical stretch on junctional AHPH (J) and junctional RhoA FRET index (K) at ZA. Data are means ± SEM from n=3 independent experiments. ****p<0.0001, ***p<0.001, **p<0.01, n.s., not significant; unpaired t-Test (D, E) with Welch’s correction or 2-way ANOVA (C, G, H, J, K) with Sidak’s multiple comparisons test. Scale bars represent 10 μm (B, G) and 60 μm (D).

We then adopted a well-used vertex-based model of an epithelial tissue to predict how these changes in cell contractility might affect cell-and tissue-level stress (Nestor-Bergmann, et al., 2017; Fletcher, et al., 2014; Farhadifar, et al., 2007; Nagai and Honda, 2001; Honda and Eguchi, 1980). We inferred patterns of relative tension and compression in the tissue using the isotropic component of the cell stress tensor, which we term the effective cell pressure, *P*^eff^ (described in the Computational Supplement). Cells satisfying *P*^eff^> 0 are predicted to be under net tension, whereas those satisfying *P*^eff^< 0 are under predicted net compression. Simulating calyculin by inducing additional contractility in the bulk and cortex of the cell (Computational Supplement), the model predicts that the proportion of cells under net tension increases following treatment with calyculin (Figure 1D and Movie S1). For a monolayer with boundary conditions that constrain its size, as might be expected of confluent epithelial monolayers, this leads to a predicted increase in global tissue pressure, but very little movement in the tissue (Movie S1).

We then examined the morphological response of monolayers using endogenous E-cadherin that was CRISPR-Cas9-engineered to bear a GFP tag at its C-terminus (Liang, et al., 2017). Consistent with predictions of the model, little movement of the cells was observed, but junctions appeared to tauten almost immediately after adding calyculin (Movie S2) and remained intact for ∼12 min, until they fractured at the very end of the movies. The junctional tautening was consistent with our recent observation that calyculin increases junctional tension, as measured by their recoil following laser ablation (Acharya, et al., 2017). We further confirmed this at the molecular level using a FRET-based tension sensor incorporated into α-catenin (αCat-TS) (Acharya, et al., 2017). Calyculin decreased energy transfer across αCat-TS (Figure 1E and S1C), consistent with an increase in tension. This was supported by increased staining for the tension-sensitive α-18 epitope of α-catenin (Figure 1F and S1D). Thus, the predicted increase in tissue tension was associated with an observed increase in tension at AJ.

We then used a location biosensor for active GTP-RhoA (AHPH, derived from the C-terminus of anillin (Piekny and Glotzer, 2008)) to monitor the response of RhoA signaling to calyculin. As previously reported (Priya, et al., 2015; Ratheesh, et al., 2012), active RhoA signaling is found in a prominent zone at the zonula adherens (ZA) in established steady-state epithelial monolayers (Figure 1G). However, the intensity of the signal in this zone increased rapidly with calyculin and remained the most prominent site for RhoA signaling in the cells (Figure 1G and 1H). By contrast, little change in GTP-RhoA was evident at cell-substrate interfaces (Figure S1E and S1F). This implied that tissue stress stimulated RhoA signaling at the ZA. This was supported with a FRET-based activity sensor that showed increased energy exchange on addition of calyculin (Figure 1I and S1G). The specificity of response for both AHPH and RhoA FRET was confirmed by inhibition with C3-transferase (C3-T; Figure 1H and 1I). Together, these findings suggested that contractile tension stimulates junctional RhoA signaling.

To reinforce this conclusion, we adopted the complementary approach of growing Caco-2 cells on flexible substrata and then subjecting them to an acute equibiaxial stretch (10%, 10 min; Figure S1H). Stretch increases the proportion of cells under net tension within a tissue (Nestor-Bergmann, et al., 2018b; Nestor-Bergmann, et al., 2017; Wyatt, et al., 2015) and we confirmed that tension on the cadherin complex increased with both αCat-TS and α-18 staining (Figure S1I to S1L). Again, we observed increased junctional RhoA signaling, measured both with AHPH (Figure 1J) and the Rho-FRET sensor (Figure 1K and S1O). Together, we conclude that junctional RhoA is a potential mechanosensitive signal that responds to tensile monolayer stress. In order to dissect its functional significance, it was then necessary to define the molecular pathway responsible for this tension-activated RhoA signaling.

### p114 RhoGEF mediates stress-activated RhoA signaling

We first sought to identify the molecule(s) responsible for activating RhoA signaling in response to stress. We screened candidate RhoA guanine nucleotide exchange factors (GEFs) that had earlier been implicated in mechanotransduction and/or junctional signaling (Figure S2A)(Lessey, et al., 2012; Ratheesh, et al., 2012; Guilluy, et al., 2011). This led us to focus on p114 RhoGEF, whose junctional localization increased clearly when monolayer stress was induced either with calyculin (Figure 2A and 2B) or stretch (Figure S2A and S2B). This was supported by fluorescence recovery after photobleaching (FRAP) turnover studies, which showed that CFP-p114 RhoGEF at junctions was stabilized by calyculin (Figure 2C). In contrast, p115 RhoGEF and LARG did not localize to junctions in Caco-2 cells either before or after stretch (Figure S2A); and while Ect2, and to a minor extent GEF H1, were found at the ZA, neither changed with stress (Figure S2A and S2B).

**Figure 2.**
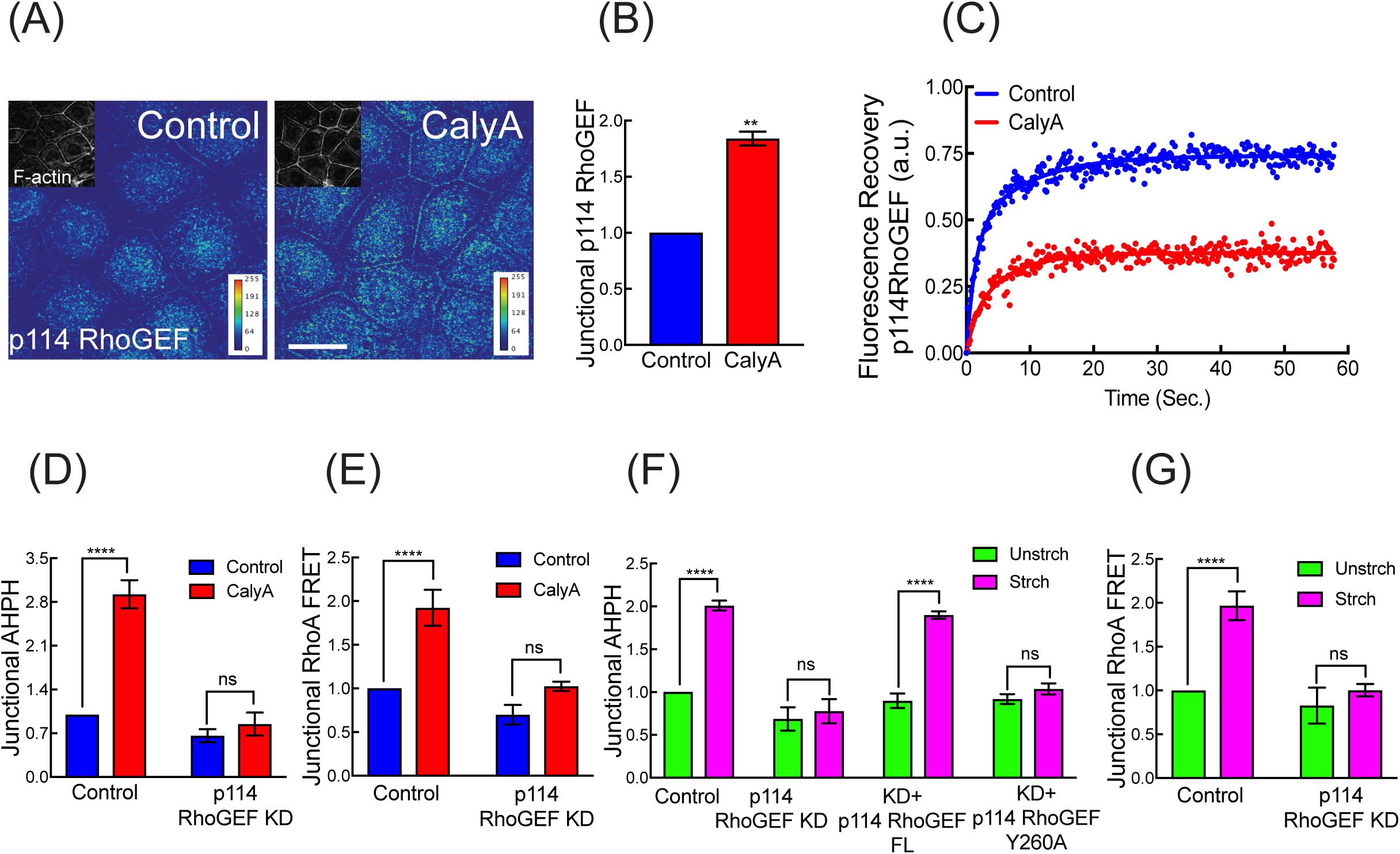
p114RhoGEF activates junctional RhoA in response to tensile stress. (A, B) Calyculin A increases active p114RhoGEF at junctions: representative images (A) and quantitation of junctional fluorescence (B). (C) Representative FRAP plot showing that calyculin A treatment increases junctional stability of CFP-p114RhoGEF^FL^. (D, E) p114RhoGEF KD inhibited calyculin A induced RhoA signaling at the ZA, measured by AHPH (D) and RhoA-FRET index (E). (F) p114RhoGEF KD inhibited stretch-induced activation of junctional RhoA, measured with AHPH. Effect of reconstitution with p114RhoGEF^FL^ and p114RhoGEF^Y260A^. (G) p114RhoGEF KD inhibited stretch-induced junctional RhoA-FRET activity. Data are means ± SEM from n=3 independent experiments. ****p<0.0001, ***p<0.001, n.s., not significant; unpaired t-Test (B) with Welch’s correction or 2-way ANOVA (D-G) with Sidak’s multiple comparisons test. Scale bars represent 20 μm (A).

We then depleted p114 RhoGEF by RNAi (knock-down, KD) to test its role in tension-activated RhoA signaling (Figure S2C). Interestingly, although p114 RhoGEF KD did not affect baseline levels of junctional AHPH robustly, it abolished the increase that was stimulated either by calyculin (Figure 2D) or stretch (Figure 2F). This was confirmed by Rho FRET (Figure 2E). In contrast, although steady-state junctional AHPH levels were decreased by Ect2 or GEF-H1 RNAi, neither affected the response to tensile stress (Figure S2D). LARG or p115 RhoGEF KD had no effect on the baseline or stretch-stimulated levels of AHPH (Figure S2D). This suggested a selective requirement for p114 RhoGEF to activate RhoA in response to tension, whereas other GEFs, such as Ect2 (Ratheesh, et al., 2012) and GEF-H1 mediated baseline signaling. The tension-activated RhoA response was restored to p114 RhoGEF KD cells by expression of an RNAi-resistant FL-p114 RhoGEF transgene (Figure 2F and S2C), confirming the specificity of this effect. However, stretch-activated RhoA signaling was not restored by expression of a GEF-defective mutant (p114 RhoGEF^Y260A^; Figure 2F and S2C). Together, these findings identified p114 RhoGEF as responsible for stimulating junctional RhoA signaling in response to monolayer stress.

### Gα12 confers junctional specificity on p114 RhoGEF recruitment

Then, we asked what was responsible for the selective junctional recruitment of p114 RhoGEF and subsequent RhoA activation. p114 RhoGEF is one of a number of GEFs that respond to heterotrimeric G-proteins of the Gα12/13 subclass (Martin, et al., 2016; Siehler, 2009; Goulimari, et al., 2005). As these G-proteins have also been implicated in cadherin-dependent morphogenesis (Kerridge, et al., 2016; Lin, et al., 2009) and can interact with the cadherin complex (Kaplan, et al., 2001; Meigs, et al., 2001), they were interesting candidates for junctional mechanotransduction. Indeed, Gα12 was faintly detectable at the ZA in unstressed monolayers but increased when stress was applied, either with calyculin (Figure 3A and S3A) or extrinsic stretch (Figure 3B and S3B). In contrast, Gα13 was scarcely detectable at junctions in these cells, either at baseline or after calyculin (Figure S3D and S3E).

**Figure 3.**
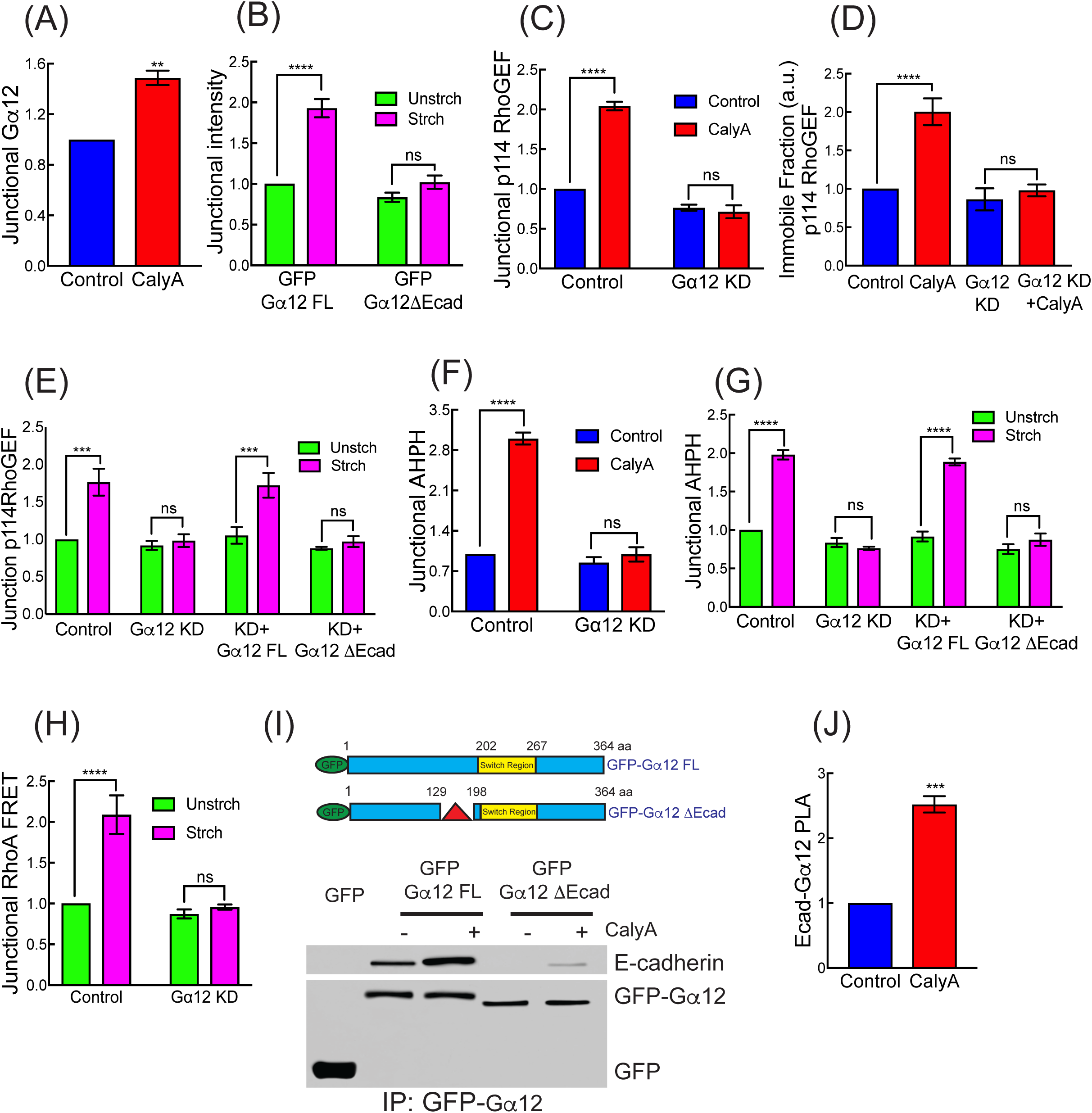
Role of heterotrimeric Gα12 in tension-activated RhoA signaling. (A) Calyculin A increases junctional Gα12. (B) Mechanical stretch increases junctional GFP-Gα12^FL^ but not GFP-Gα12^ΔEcad^ reconstituted in Gα12 RNAi cells. (C, D) Gα12 siRNA reduces junctional p114RhoGEF (C) and stabilization CFP-p114RhoGEF (D, immobile fraction in FRAP) in response to calyculin A. (E) Effect of Gα12 siRNA and reconstitution with GFP-Gα12^FL^ or GFP-Gα12^ΔEcad^ on junctional recruitment of p114RhoGEF in response to mechanical stretch. (F, G) Effect of Gα12 siRNA (KD; F,G) and reconstitution with GFP-Gα12^FL^ or GFP-Gα12^ΔEcad^ (G) on tension-activated junctional RhoA signaling measured by AHPH in response calyculinA (F) and mechanical stretch (G). (H) Gα12 siRNA reduces junctional RhoA-FRET index in mechanically stretched cells. (I) Schematic representation of GFP-Gα12^FL^ and GFP-Gα12^ΔEcad^ constructs (Δ showing the deletion site on Gα12ΔEcad) on top. Bottom, GFP-Gα12^FL^ or GFP-Gα12^ΔEcad^ expressed in Gα12 were isolated by GFP-Trap and immunoblotted for E-cadherin. (J) Calyculin A increases PLA reaction between junctional E-cadherin and Gα12. Data are means ± SEM from n=3 independent experiments. ****p<0.0001, ***p<0.001, **p<0.01, n.s., not significant; unpaired t-Test (A, J) with Welch’s correction or 2-way ANOVA (B-H) with Sidak’s multiple comparisons test. Scale bars represent 20 μm (J).

We further found that junctional recruitment of p114 RhoGEF by calyculin (Figure 3C) or stretch (Figure 3E) was impaired in Gα12 RNAi cells (Figure S3C). Stretch-induced recruitment of p114 RhoGEF was restored to Gα12 RNAi cells by expression of an RNAi-resistant FL-Gα12 transgene (Figure 3E). In contrast, stretch-induced recruitment of p114 RhoGEF was not affected by Gα13 KD (Figure S3G). This suggested that Gα12 was selectively responsible for recruiting p114 RhoGEF to junctions in response to tensile stress. This was reinforced by FRAP experiments, which showed that calyculin did not stabilize CFP-p114 RhoGEF in Gα12 RNAi cells (Figure 3D and S3F). Importantly, neither calyculin (Figure 3F) nor mechanical stretch (Figure 3G) increased junctional RhoA signaling measured by AHPH when Gα12 was depleted unlike Gα13 KD (Figure S3H). The specificity of this effect was established by rescue with an RNAi-resistant FL-Gα12 transgene (Figure 3G). The Rho FRET sensor confirmed that Gα12 was necessary for stretch to activate junctional RhoA (Figure 3H). Therefore, Gα12 was necessary for tensile stress to activate junctional RhoA signaling via p114 RhoGEF. As had been observed for p114 RhoGEF, Gα12 KD did not materially affect the baseline level of junctional RhoA activity, but selectively perturbed the response to tensile stress (Figure 3F and 3G).

Earlier studies reported that Gα12 can bind directly to the C-terminus of E-cadherin (Kaplan, et al., 2001; Meigs, et al., 2001). We therefore hypothesized that this interaction might confer junctional selectivity on the tension-activated RhoA response. To test this, we expressed GFP-tagged Gα12 in Gα12 RNAi cells and isolated protein complexes with GFP-Trap (Figure 3I). E-cadherin associated with GFP-Gα12 at baseline and the amount was substantially increased by calyculin (Figure 3I). This was supported by proximity ligation analysis (PLA), which showed an increased reaction between the two endogenous proteins at cell-cell junctions when tension was applied to monolayers (Figure 3J and S3I). We then deleted a 67-amino acid region in Gα12 implicated in binding E-cadherin, which is separate from the switch region that activates p114 RhoGEF (Gα12^ΔE-cad^, Figure 3I). Gα12^ΔE-cad^ behaved as a cadherin-uncoupled mutant, as it did not co-immunoprecipitate with E-cadherin at baseline and only a trace increase was seen after stimulation with calyculin (Figure 3I). Nor was Gα12^ΔE-cad^ recruited to junctions when monolayers were stretched (Figure 3B). Importantly, Gα12^ΔE-cad^ reconstituted in Gα12 KD cells did not support either the tension-sensitive junctional recruitment of p114 RhoGEF (Figure 3E) or the stretch-stimulated increase in junctional GTP-RhoA (Figure 3G). Together, these findings indicate that tensile stress activates RhoA signaling at the ZA by recruiting Gα12 to E-cadherin. Gα12 then represented a key intermediate that allowed force-sensing at AJ to be transduced into chemical signaling by RhoA.

### Myosin VI recruits to E-cadherin in response to tensile stress

The selective junctional localization of GTP-RhoA and its activators suggested that AJ might possess a mechanosensor responsible for detecting acute increases in tensile stress. We focused our attention on Myosin VI, an F-actin-binding motor that can couple adhesion to the cytoskeleton by directly binding E-cadherin (Mangold, et al., 2012; Maddugoda, et al., 2007). Importantly, Myosin VI has an intrinsic force-sensitivity (Chuan, et al., 2011; Oguchi, et al., 2008; Altman, et al., 2004). Under low loads, dimeric Myosin VI serves as a processive motor, but it can convert to a dynamic F-actin-based anchor when sufficient load is applied (Chuan, et al., 2011). We hypothesized that this property might allow Myosin VI to sense tensile forces at the ZA. Consistent with a potential role in force-sensing, junctional Myosin VI levels were increased when monolayer stress was applied (Figure 4A to 4C and S4A) and FRAP studies showed that calyculin stabilized Myosin VI-GFP at the ZA (Figure 4D, S4B and S4C). Calyculin also increased the amount of Myosin VI that co-precipitated with E-cadherin-GFP (Figure 4E, 4F and S4D), as well as the interaction between the endogenous proteins detected by PLA (Figure 4G and S4E). Thus, monolayer stress promoted the association with E-cadherin that recruits Myosin VI to junctions (Maddugoda, et al., 2007).

**Figure 4.**
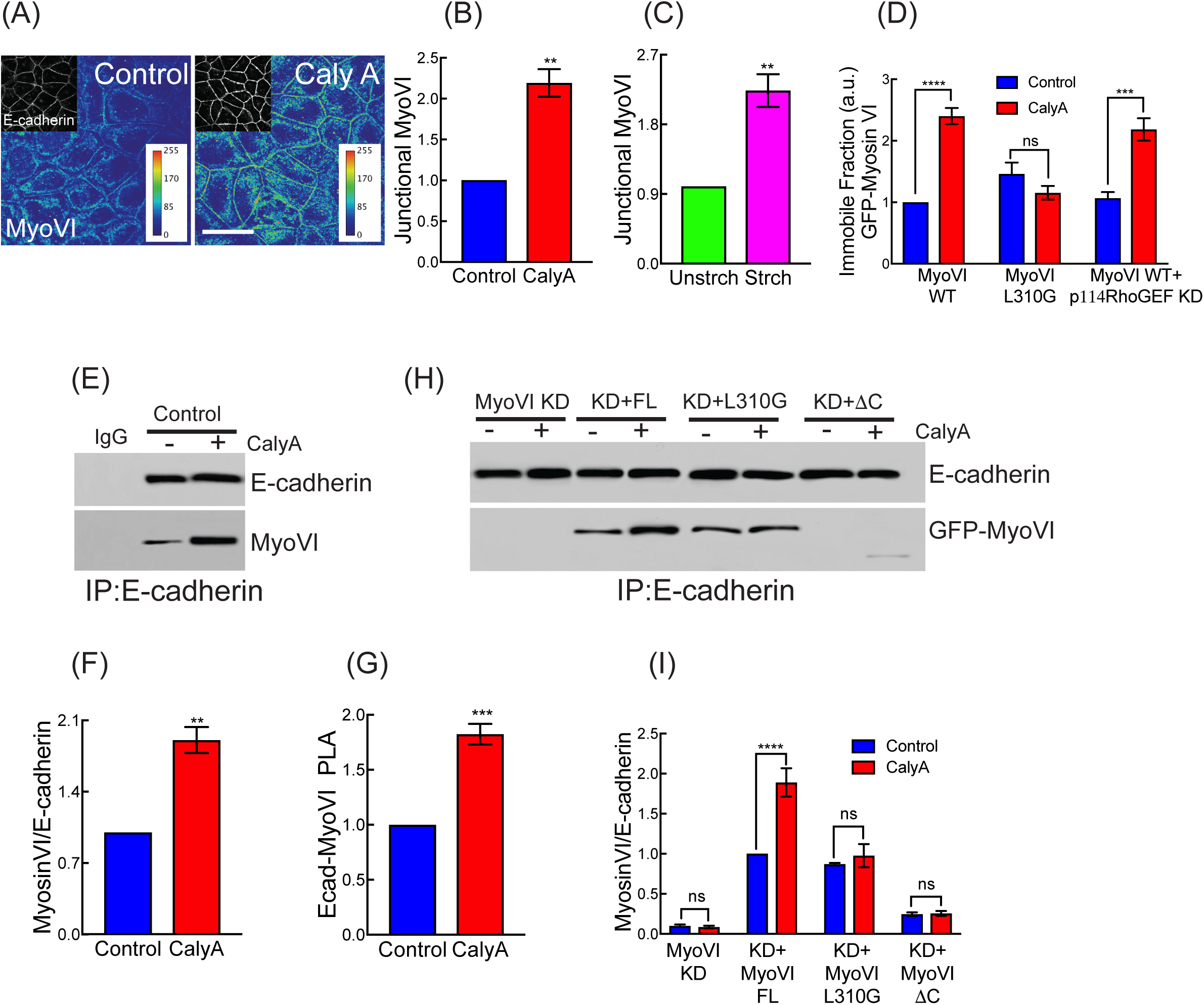
*Myosin VI recruits to E-cadherin in response to tensile monolayer stress.* (A-C) Effect of Calyculin A (A,B) and mechanical stretch on junctional myosin VI (C): representative images (A) and quantitation (B, C). (D) Junctional stability measured by FRAP of GFP-Myosin VI^WT^ or GFP-Myosin VI^L310G^. Stabilization of GFP-Myosin VI^WT^ is not affected by p114 RhoGEF RNAi. (E, F) Effect of calyculin A on co-Immunoprecipitation of endogenous Myosin VI with endogenous E-cadherin: representative blot (E) and quantitation (F). (G) Calyculin A increases the PLA reaction between junctional E-cadherin and Myosin VI. (H, I) Effect of calyculin A on co-immunoprecipitation of E-cadherin with GFP-Myosin VI constructs, expressed in Myosin VI RNAi cells: representative blot (H) and quantitation (I). Data are means ± SEM from n=3 independent experiments. ****p<0.0001, ***p<0.001, **p<0.01, n.s., not significant; unpaired t-Test (B, C, F, G) with Welch’s correction or 2-way ANOVA (D, I) with Sidak’s multiple comparisons test. Scale bars represent 20 μm (A).

To test whether this might involve the intrinsic force sensitivity of Myosin VI, we sought to specifically disrupt force sensitivity while retaining the transport function of the motor. Force sensitivity in Myosin VI is thought to arise from mechanical gating of nucleotide exchange. Applied load has been reported to accelerate ADP binding, driving a transition from transport to anchoring in dimers (Chuan, et al., 2011; Altman, et al., 2004). Force-dependent inhibition of ADP release (Elting, et al., 2011; Dunn, et al., 2010) or acceleration of ATP binding (Sweeney, et al., 2007) have also been proposed as mechanisms for coordinating heads via intramolecular strain. The insert 1 region of the motor is implicated in nucleotide gating, and a point mutant in this region (Myosin VI ^L310G^) affects nucleotide exchange and abolishes ATPase gating in the dimer (Pylypenko, et al., 2011). However, this mutant preserves processive transport function, with only minor reductions in processive run length (Pylypenko, et al., 2015).

We therefore hypothesized that the L310G mutation would specifically affect load-dependent anchoring by Myosin VI *in vivo.* Then we reconstituted Myosin VI KD cells with Myosin VI^L310G^ to test if this mutation affected its tension-sensitive recruitment (Figure S4F). We found that, in contrast to GFP-Myosin VI^FL^, GFP-Myosin VI^L310G^ was not recruited to junctions (Figure S4A) or stabilized in response to calyculin (Figure 4D, S4B and S4C). We further compared the biochemical association of these transgenes with E-cadherin (Figure 4H and 4I). As seen for the endogenous protein, calyculin increased the association of GFP-Myosin VI^FL^ with E-cadherin, and this required the cargo-binding domain that binds E-cadherin (Myosin VI^ΔC^) (Mangold, et al., 2012). In contrast, while GFP-Myosin VI^L310G^ interacted with E-cadherin under basal conditions, its association was not increased in response to calyculin (Figure 4H and 4I). Therefore, processive motor function, which is retained in Myosin VI^L310G^, is not sufficient for stress-induced junctional recruitment. Instead, these findings suggest that recruitment of Myosin VI reflects its load-sensitive anchorage mediated by nucleotide gating.

### Myosin VI mediates tension-activated RhoA signaling

Interestingly, junctional recruitment of Myosin VI preceded the accumulation of AHPH when cells were stimulated with calyculin (Figure S5A). This suggested that Myosin VI might be an upstream element in the RhoA-activation pathway. Indeed, Myosin VI RNAi abolished the stretch-induced increase in junctional RhoA, monitored with either the AHPH (Figure 5A and 5B) or Rho FRET sensors (Figure 5C and S5B). Consistent with its acting at an upstream point in tension-activated RhoA signaling, Myosin VI KD also compromised the stretch-induced recruitment of Gα12 (Figure 5D and 5E) and p114 RhoGEF (Figure S5C and S5D). In contrast, p114 RhoGEF KD did not affect tension-sensitive recruitment of Myosin VI (Figure S5E). All these features were restored by Myosin VI ^FL^ but not by GFP-Myosin VI^L310G^, implying that its intrinsic mechanosensitivity was necessary for Myosin VI to participate in tension-activated RhoA signaling.

**Figure 5.**
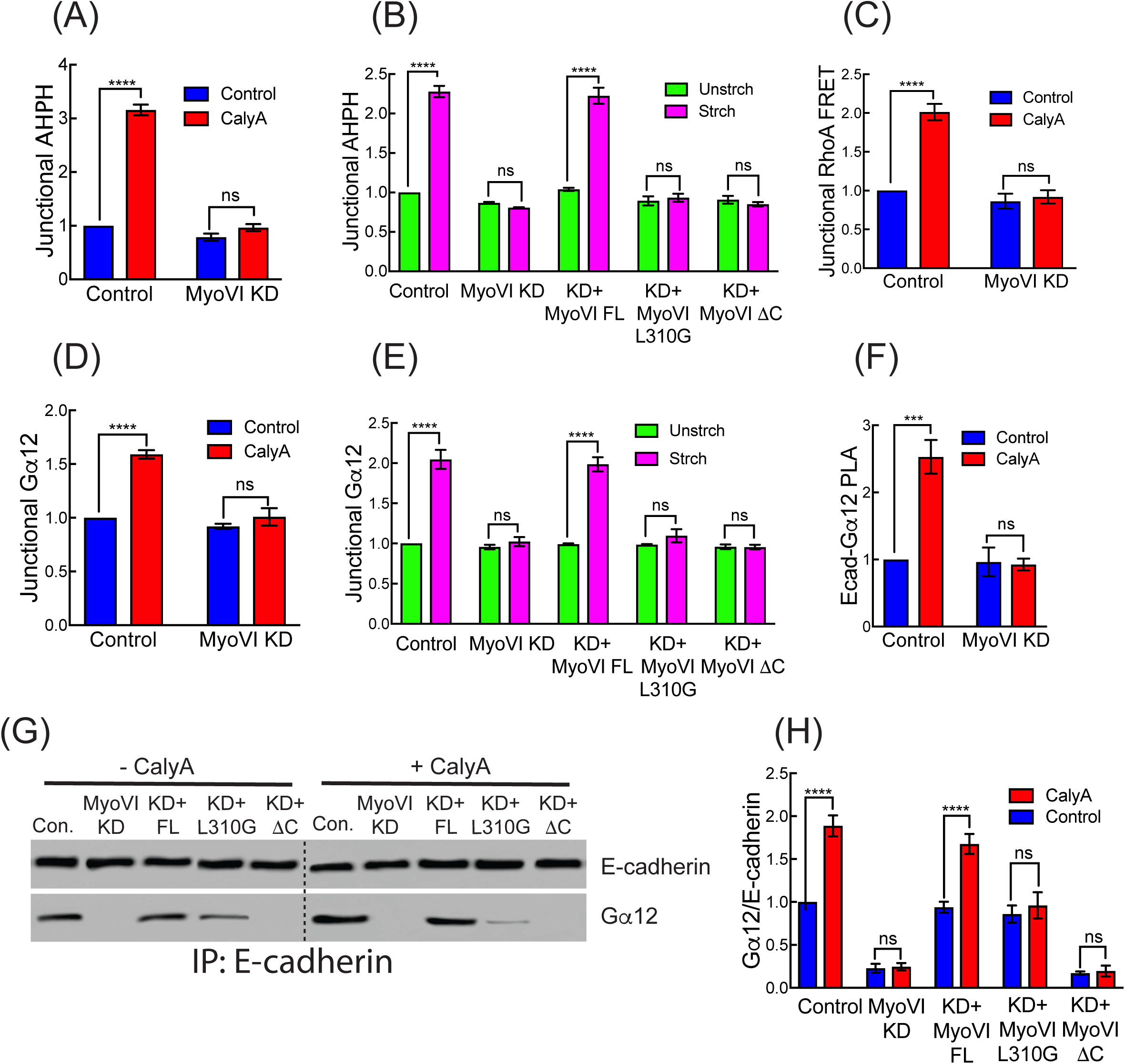
Myosin VI supports tension-activated junctional RhoA signaling. (A) Effect of Myosin VI KD on tension-activated RhoA signaling, measured by AHPH in calyculin A treated cells. (B) Effect of Myosin VI mutations on tension-activated RhoA signaling, measured by AHPH in stretch-stimulated monolayers. Myosin VI RNAi cells were reconstituted with GFP-Myosin VI^FL^, GFP-Myosin VI^L310G^ or GFP-Myosin VI^ΔC^. (C) Myosin VI KD reduces calyculin-activated junctional RhoA signaling measured by RhoA-FRET index. (D, E) Effect of Myosin VI RNAi (D,E) and reconstitution with Myosin VI transgenes (E) on junctional recruitment of Gα12 in response to calyculin A (D) or mechanical stretch (E). (F) Myosin VI KD reduces the calyculin-induced PLA reaction between E-cadherin and Gα12. (G, H) Effect of Myosin VI mutations on the calyculin-stimulated association of GFP-Myosin VI and E-cadherin; transgenes were expressed in Myosin VI RNAi cells: representative blot (G) and quantitation (H). Data are means ± SEM from n=3 independent experiments. ****p<0.0001, ***p<0.001, **p<0.01, n.s., not significant; 2-way ANOVA (A-F and H) with Sidak’s multiple comparisons test.

How, then, did Myosin VI signal to recruit the Gα12-p114 RhoGEF apparatus? Since Myosin VI can interact with E-cadherin, we considered the hypothesis that it might promote the association between E-cadherin and Gα12. Indeed, we found that an increased amount of Gα12 co-immunoprecipitated with E-cadherin when monolayers were treated with calyculin and this was reduced by Myosin VI KD (Figure 5G, 5H and S5F). Similarly, Myosin VI KD reduced the calyculin-stimulated association between Gα12 and E-cadherin that was detected at junctions by PLA (Figure 5F). The biochemical interaction between E-cadherin and Gα12 was restored by expression of Myosin VI^FL^, but not when cells were reconstituted with either Myosin VI^ΔC^ or Myosin VI^L310G^ (Figure 5G and 5H). Thus, both the intrinsic load sensitivity of Myosin VI and its ability to bind E-cadherin were necessary for tensile stress to enhance the E-cadherin-Gα12 interaction. As Myosin VI was also found in the E-cadherin immune complexes, it was formally possible that Myosin VI might independently recruit Gα12. However, the association between Myosin VI and Gα12 required E-cadherin, being reduced E-cadherin KD (Figure S5G to S5I). This suggested that Myosin VI interacts indirectly with Gα12 through E-cadherin. Overall, these data identify Myosin VI as the mechanism that couples force sensing to signal transduction by promoting the formation of an E-cadherin-Gα12 complex that leads to activation of RhoA.

### p114 RhoGEF signaling helps monolayers resist tensile stress

Altogether, these findings identified a mechanosensitive RhoA pathway at AJ that responds when tensile stress is applied to monolayers. To evaluate its functional significance, we first tested how the epithelial barrier, measured by transepithelial electrical resistance (TER), responded when intrinsic tension was increased with calyculin. TER was effectively preserved in control monolayers, even when they were treated with calyculin (Figure 6A and S6A). Similarly, p114 RhoGEF KD cells showed a stable TER throughout the observation period (Figure 6A). However, TER rapidly and progressively fell when calyculin was added to p114 RhoGEF KD cells (Figure 6A and S6A). The implication that RhoA signaling was necessary to preserve the epithelial barrier in the face of tensile stress was further supported by blocking RhoA directly with C3-T (Figure 6A and S6A). Together, these findings suggested that the stress-activated RhoA pathway was necessary to preserve epithelial barrier integrity in the face of monolayer stress.

**Figure 6.**
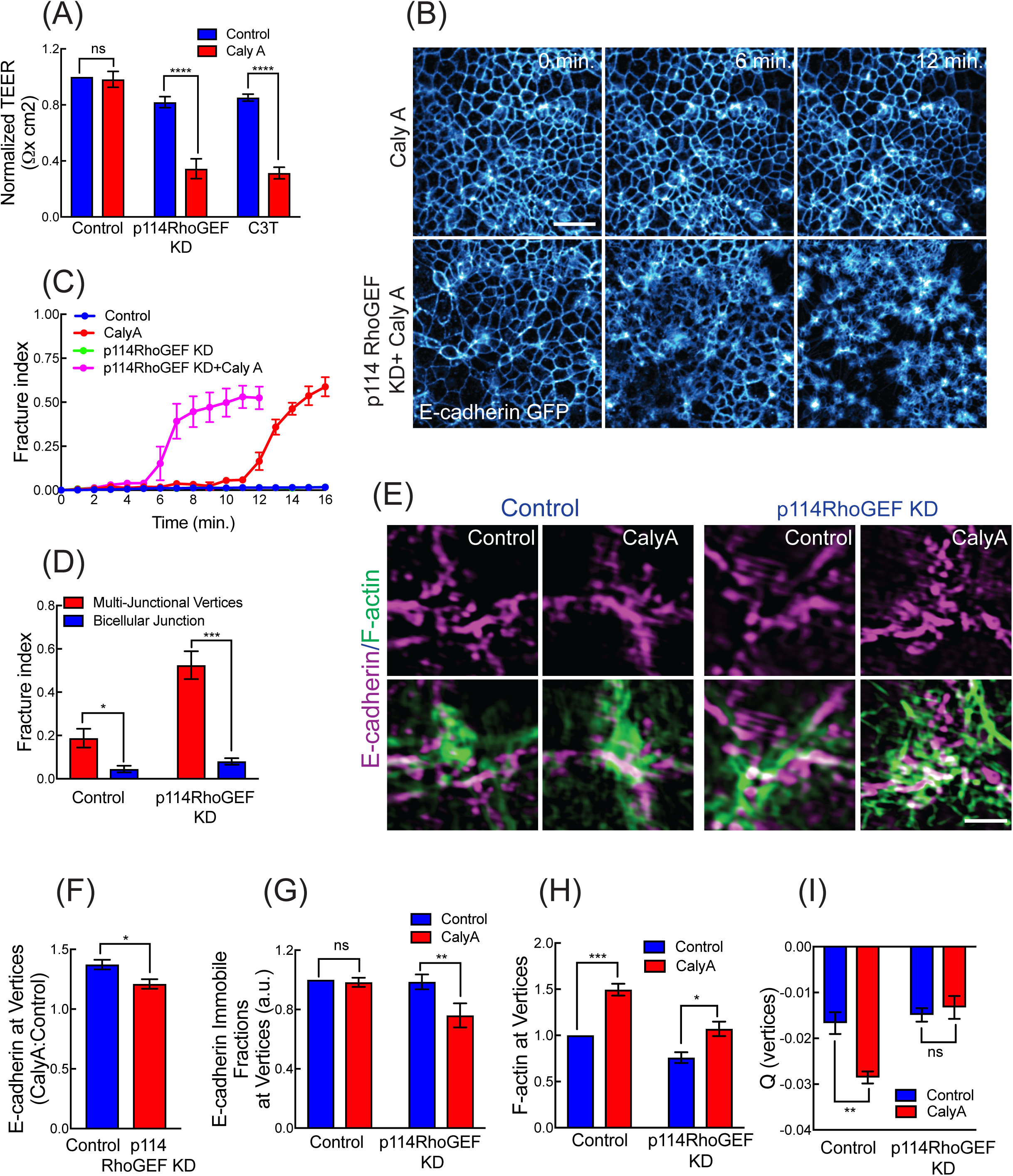
Tension-activated junctional RhoA signaling preserves epithelial integrity. (A) Effect of p114RhoGEF KD or C3T on epithelial transepithelial electrical resistance (TER) after stimulation with calcyulin A. Monolayers were grown for 21 days, pre-treated with 3CT as appropriate, then stimulated with calyculin for 15 min. (B, C) Effect of p114 RhoGEF KD on sensitivity of monolayers to fracture following calyculin: representative stills from E-cadherin-GFP movies (B, corresponding movies are Movie 2 and Movie 5) and quantitation (C). (D) Calyculin A triggered monolayer fracture preferentially occurs at multicellular vertices. (E) E-cadherin and F-actin immunofluorescence at multicellular vertices imaged by SIM. (F) Effect of p114RhoGEF KD on calyculin-induced accumulation of E-cadherin at multicellular vertices. Data are ratios of E-cadherin fluorescence intensity in calyculin-stimulated compared with control culturs. (G) Effect of p114RhoGEF KD on immobile fraction of E-cadherin at multicellular vertices. (H, I) Effect of p114RhoGEF KD on fluorescence intensity (H) and nematic order (I) of F-actin at multicellular cell vertices in control and calyculin-stimulated cells. (I) p114RhoGEF KD increases Nematic order of F-actin filament orientation at Data are means ± SEM from n=3 independent experiments. ****p<0.0001, ***p<0.001, **p<0.01, *p<0.05, n.s., not significant; unpaired t-Test (F) with Welch’s correction or 2-way ANOVA (A, D and G-I) with Sidak’s multiple comparisons test. Scale bars represent 60 μm (B) and 2 μm (H).

To better characterize this process, we monitored E-cad-GFP junctions by live-cell imaging. Monolayer integrity remained undisturbed in both control Caco-2 cells (Movie S3) and p114 RhoGEF KD cells (Movie S4) that were not treated wit calyculin. In contrast, calyculin caused p114 RhoGEF KD cells to fracture at multiple sites throughout the monolayer during the course of the movies (Movie S6). Although fracturing was eventually seen in control monolayers treated with calyculin (Movie S2), quantitation confirmed that it occurred much earlier in p114 RhoGEF KD cells (Figure 6B and 6C). Collectively, these findings indicated that the p114 RhoGEF pathway serves to preserve the integrity of cell-cell contacts against the increased tensile stress within the monolayers.

### Mechanical modelling predicts that p114 RhoGEF signaling protects epithelial integrity by increasing yield limit

We then used the vertex model to consider how tension-activated p114 RhoGEF signaling might alter cell mechanics to maintain epithelial integrity (Computational Supplement). One possibility was that p114 RhoGEF signaling increased the stiffness of the junctional cortex, which was predicted to protect a tissue undergoing a stretch deformation by decreasing the bulk modulus while increasing the shear modulus of the tissue (Nestor-Bergmann, et al., 2018a). Actomyosin contributes to cortical stiffness and, indeed, junctional actomyosin increased when control monolayers were treated by calyculin, but not in p114 RhoGEF KD cells (Figure S6B and S6C). However, when we modelled an increase in internal contractility induced by calyculin, simulations revealed that further increasing cortical stiffness alone accelerated the onset of fractures (Movie S7), rather than delaying it, as was observed experimentally (Figure 6C). Thus, an increase in cortical stiffness alone did not explain the protective effect of p114 RhoGEF signaling.

To obtain further insight into the additional protective effect of the p114 pathway, we first assessed where fractures were initiated. Physical considerations predicted that multicellular vertices would be the sites that were subject to the greatest level of force (Higashi and Miller, 2017) and therefore the most prone to fracture. Indeed, quantitation of E-Cad-GFP movies confirmed that calyculin-induced fractures began overwhelmingly at vertices (Figure 6D), in both control and p114 RhoGEF KD cells. We therefore hypothesized that an increase in the yield limit of multicellular vertices might protect monolayers, even when cortical stiffness was also increased. Supporting this, simulations showed that increasing the yield limit substantially delayed the onset of fracture (Movie S8), as was found in our experiments. Furthermore, the fracture behaviour resembled more closely that seen experimentally. Thus, the increased build-up of forces led to cell recoil after fracture that was much greater than that seen in p114 RhoGEF KD monolayers (compare Movies S2 and S8 with Movie S5).

Close inspection of the movies indicated that cells separated from one another as fractures began (Figure S6D, Movies S9 and Movie S10), suggesting a defect in adhesion. Indeed, E-cadherin intensity increased at vertices when control monolayers were treated with calyculin, but did not rise in p114 RhoGEF KD cells (Figure 6E, F). This was further supported by FRAP analyses, which showed that the immobile fraction of E-cadherin-GFP was reduced when p114 RhoGEF KD cells were treated with calyculin, whereas it was maintained in control cells (Figure 6G).

We then examined the actomyosin cytoskeleton at the vertices, as a potential locus for cadherin stabilization. Since NMII stimulation by calyculin was not materially affected by p114 RhoGEF KD (Figure S6E and SF), we focused on F-actin itself. The F-actin content at the vertices was increased when control cells were stimulated by calyculin (Figure 6G) and SIM imaging showed that this manifested as a dense accumulation (Figure 6E), with increased nematic order (Figure 6I), suggesting that F-actin organization was becoming more co-linear. In contrast, F-actin organization at vertices was more disorganized at baseline in p114 RhoGEF KD cells (Figure 6E), and failed to condense (Figure 6H) or increase in nematic order after calyculin (Figure 6I). Overall, these findings suggest that mechanosensitive p114 RhoGEF signaling to the junctional cytoskeleton may help maintain a stable fraction of E-cadherin at vertices that is necessary for monolayers to resist tensile stresses.

## Discussion

Epithelia are subject to tensile forces that can potentially disrupt the cell-cell integrity that is necessary for them to form physiological barriers (Charras and Yap, 2018). This is exemplified by our observation that monolayers fracture through separation of cell-cell contacts when monolayer contractility is acutely increased by calyculin. Our experiments now identify a junctional mechanotransduction pathway that is responsible for sensing, and responding to, such tensile stresses. We propose that Myosin VI is the key sensor of tensile stress applied to adherens junctions that promotes the formation of an E-cadherin-Gα12 complex to activate the p114 RhoGEF-RhoA pathway (Figures S6H, S7). Of note, RhoA signaling is active at the ZA, even under resting conditions (Ratheesh, et al., 2012), but this is mediated by distinct GEFs, such as Ect2 and GEF-H1. Thus, the Myosin VI-Gα12-p114 RhoGEF pathway that we have identified can be considered a selective response to suprabasal tensile stress.

At first sight, it seemed paradoxical that stimulation of RhoA would be used to preserve epithelial integrity. RhoA promotes actomyosin assembly at AJ under resting conditions (Ratheesh, et al., 2012; Smutny, et al., 2010) and also in calyculin-stimulated cells. Both F-actin and NMII increased at bicellular junctions upon treatment with calyculin and this was abrogated by p114 RhoGEF KD. Although RhoA can activate NMII and also promote actin assembly (Lessey, et al., 2012), we consider that actin regulation was the key factor in our experiments, as calyculin appeared to maximally stimulate NMII. Nonetheless, this p114 RhoGEF-stimulated increase in actomyosin might be expected to increase the line tension in bicellular junctions and enhance the forces acting to disrupt epithelial integrity, especially when focused on multicellular junctions (Higashi and Miller, 2017). One possibility was that enhanced actomyosin also increased the stiffness of junctions, providing increased resistance to tensile stress. However, simulations in a mechanical model indicated that this increasing stiffness alone accelerated monolayer fracture rather than retarding it.

Instead, we consider that the protective effect of the p114 RhoGEF pathway is better explained by an increase in the yield limit of cell vertices. In simulations of our vertex model, increasing the yield limit protected monolayer integrity against calyculin-induced stresses, even if junctional stiffness was also increased. Experimentally, we suggest that this reflects an increase in resistance at the multicellular vertices. Physical considerations identify vertices as the junctional sites where cellular forces will be greatest (Higashi and Miller, 2017) and, indeed, vertices were the principal sites where cell separation first began in our experiments. The accelerated onset of fracture that was seen in p114 RhoGEF KD cells then implied that tension-activated p114 RhoGEF-RhoA signaling might reinforce vertices against stress. A diverse variety of membrane proteins have been identified at multicellular junctions (Higashi and Miller, 2017), including proteins that are restricted to the vertices as well as others, such as E-cadherin itself, that are found more generally. Without excluding possible contributions of other adhesion mechanisms, our data implicate a role for regulation of E-cadherin itself. Thus, E-cadherin levels increased at vertices of calyculin-stimulated cells, but this did not occur when p114 RhoGEF was depleted. Turnover studies suggest that this reflects a failure to stabilize E-cadherin, which may be due to a concomitant failure of F-actin dynamics to respond to the stress. This is consistent with earlier evidence that RhoA stimulates actin regulators, such as mDia1, which promote filament assembly and regulate F-actin network organization to stabilize E-cadherin at the ZA (Acharya, et al., 2017; Rao and Zaidel-Bar, 2016; Carramusa, et al., 2007).

It was noteworthy that RhoA signaling was selectively increased at cell-cell junctions, but not at other adhesive sites, especially cell-substrate interactions. This highlights a key role for force-sensors to confer spatial specificity on the mechanotransduction response. Here, we find that this crucial role is played by Myosin VI, which was recruited to junctions stimulated by tensile stress and was responsible for eliciting the downstream RhoA response. We suggest that this reflects the pronounced capacity of Myosin VI to anchor to actin filaments in response to load (Chuan, et al., 2011; Altman, et al., 2004), as the impact of Myosin VI was abrogated by the L310G mutant which retains processive motor function but has defective nucleotide gating linked to load-sensitivity (Pylypenko, et al., 2015; Pylypenko, et al., 2011). We propose that monolayer tension is transmitted to Myosin VI via the E-cadherin with which it associates, even under resting conditions (Maddugoda, et al., 2007). Increased load then stabilizes Myosin VI at the ZA by increasing its anchorage to the junctional F-actin cytoskeleton. Exactly how Myosin VI promotes the association of Gα12 with E-cadherin to activate RhoA is an important question for future work. One possibility is that it may promote conformational changes or recruit associated proteins that stabilize the cadherin-Gα12 interaction; alternatively, actin-anchored Myosin VI may constitute a kinetic trap that regulates cadherin clustering for signaling. Irrespective, Myosin VI appears to exert its signaling effects via E-cadherin-Gα12, since stress-activated RhoA was abolished if Gα12 was unable to bind cadherin.

In conclusion, our findings identify a mechanotransduction pathway that is selectively elicited to preserve epithelial integrity in response to tensile stress. The selectivity of this pathway implies that junctions may possess multiple mechanisms that sense mechanical signals that may be elicited under different circumstances. Of note, α-catenin is necessary for the elemental force-sensitive association of cadherins with F-actin (Buckley, et al., 2014), and also supports Ect 2-dependent RhoA signaling at junctions under basal conditions (Ratheesh, et al., 2012). Therefore, α-catenin may confer mechanosensitivity under basal conditions, whereas the Myosin VI-dependent pathway that we have identified is elicited in response to superadded stress. Furthermore, our experiments tested the effects of acute application of tensile stress. Other mechanisms can be elicited when stresses are applied more slowly or sustained longer, such as cellular rearrangements and oriented cell division (Hart, et al., 2017; Etournay, et al., 2015; Wyatt, et al., 2015; Campinho, et al., 2013). That epithelia possess such a diversity of compensatory mechanisms attests to the fundamental challenge of mechanical stress in epithelial biology.

## Acknowledgements

We thank all our laboratory colleagues for their support and advice, and the many colleagues who provided reagents. This work was supported by grants (1037320, 1067405, 19929) and fellowships (1044041, 1136592) from the National Health and Medical Research Council of Australia, Australian Research Council (DP150101367), Queensland Cancer Council (1086857, 1128123) and Human Frontiers Science Program (RGP0023/2014). Optical microscopy was performed at the ACRF/IMB Cancer Biology Imaging Facility, established with the generous support of the Australian Cancer Research Foundation and the Queensland Brain Institute microscopy facility supported by ARC LIEF grant (LE130100078). AN-B was supported by a Company of Biologists, Journal of Cell Science, Travelling Fellowship (JCSTF-170805).

## Supplementary Figure Captions

**Figure S1 related to Figure 1.**
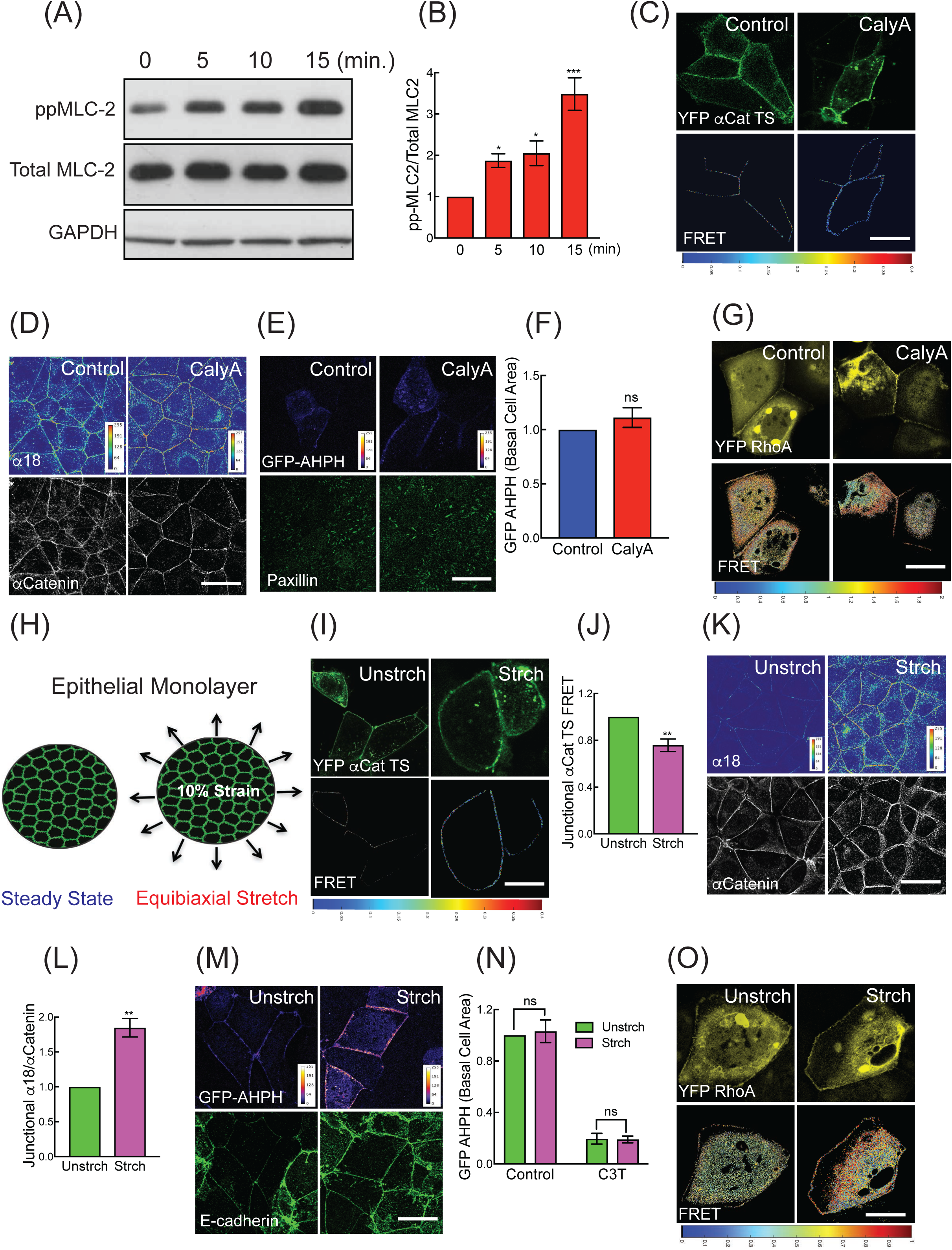
Tensile monolayer stress activates RhoA at the zonula adherens. (A, B) Activation of Myosin II (pp-MLC2, pT18/S19) in response to calyculin: representative blot (A) and quantitation (B) (C, D) Representative images of junctional αE-Cat-TS FRET (C) and junctional α18 (D) in calyculin A treated cells. (E, F) Effect of calyculin A on Basal RhoA signaling measured by AHPH: representative images (F) and quantitation. (G) Representative images of junctional RhoA-FRET in calyculin A treated cells. (H) Cartoon of equibiaxial mechanical stretching of Caco-2 monolayers grown on an elastic membrane (10% static strain, radial=circumferential). (I,J) Mechanical stretch decreases junctional αE-Cat-TS FRET: representative images (I) and quantitation (J). (K-L) Mechanical stretch increases junctional α18: representative images (K) and quantitation (L). (M) Representative images of junctional AHPH upon mechanical stretch. (N) Effect of mechanical stretch on basal AHPH: quantitation. (O) Representative images of junctional RhoA-FRET in mechanically stretched cells. Data are means ± SEM from n=3 independent experiments. ***p<0.001, **p<0.01, *p<0.05, n.s., not significant; unpaired t-Test (F, H, L) with Welch’s correction, One way Anova (B) with Dunnett’s post hoc test or 2-way ANOVA (J) with Sidak’s multiple comparisons test. Scale bars represent 10 μm (C, E, G, I, M, O) or 20 μm (D, K).

**Figure S2 related to Figure 2.**
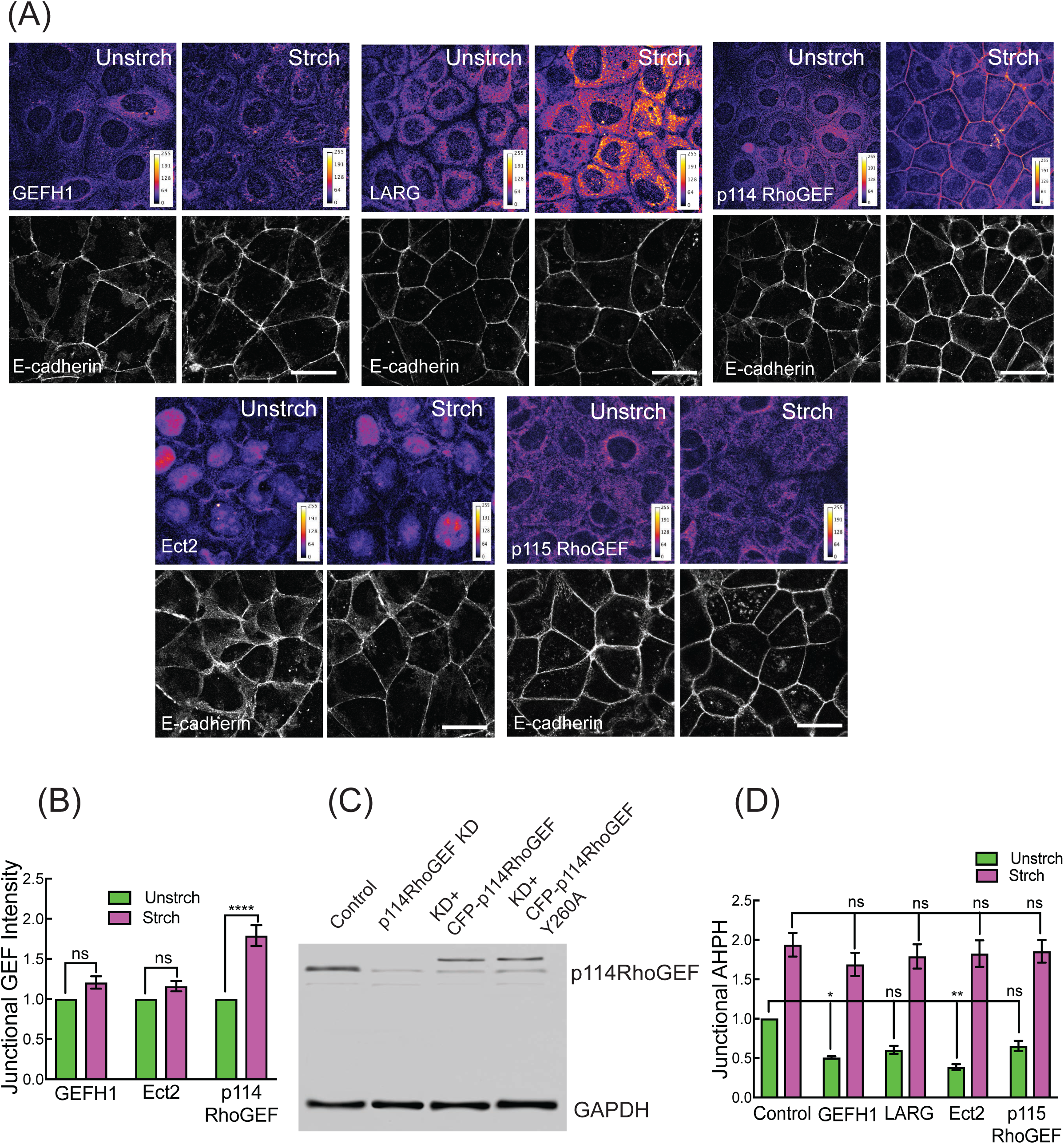
p114RhoGEF activates junctional RhoA in response to tensile stress. (A, B) Effect of mechanical stretch on different RhoA GEFs at ZA. Representative intensity images (A) and quantitation (B). (C) Representative immunoblot showing p114 RhoGEF KD and reconstitution with CFP-p114 RhoGEF^WT^ and p114 RhoGEF^Y260A^. (D) Effect of GEF RNAi on stretch-activated junctional RhoA signaling measured by AHPH. Data are means ± SEM from n=3 independent experiments. ****p<0.0001, **p<0.01, *p<0.05, n.s., not significant; 2-way ANOVA (B, D) with Sidak’s multiple comparisons test. Scale bars represent 20 μm (A).

**Figure S3 related to Figure 3.**
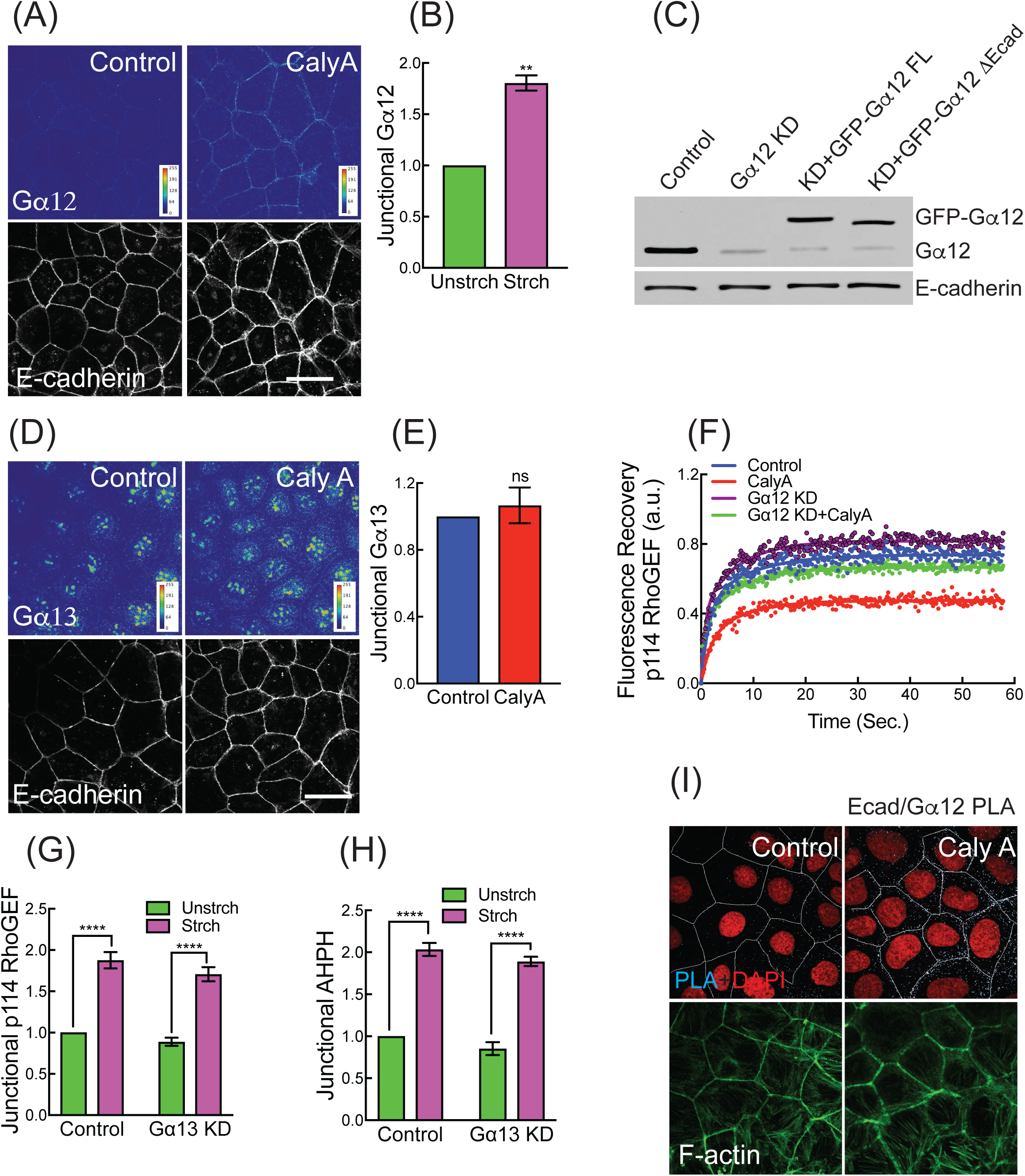
Role of heterotrimeric Gα12 in tension-activated RhoA signaling. (A, B) Calyculin A increases junctional Gα12: representative images (A) and quantitation (B). (C) Representative immunoblot of Gα12 KD and reconstitution with GFP-Gα12^WT^ and Gα12^ΔEcad^. (D, E) Junctional Gα13 is unaffected by calyculin A: representative images (C) and quantitation (D). (F) Effect of Gα12 KD on FRAP of CFP-p114RhoGEF in calyculin A treated cells. (G, H) Junctional p114RhoGEF (F) and junctional AHPH (G) is unaffected by Gα13 KD in mechanically stretched cells. Data are means ± SEM from n=3 independent experiments. ****p<0.0001, **p<0.01, n.s., not significant; unpaired t-Test (B, E) with Welch’s correction or 2-way ANOVA (F, G) with Sidak’s multiple comparisons test. Scale bars represent 20 μm (G, H).

**Figure S4 related to Figure 4.**
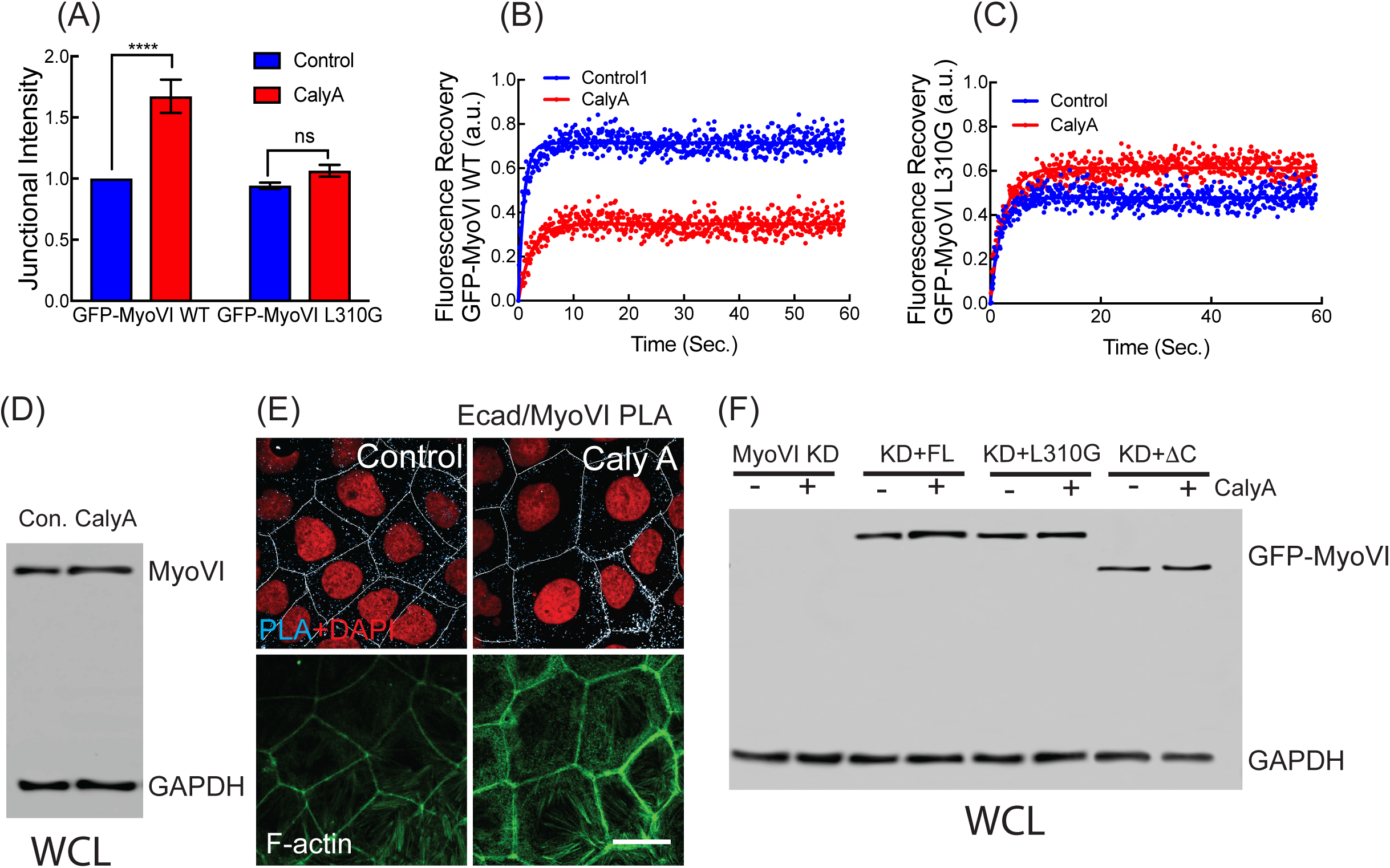
Myosin VI recruits to E-cadherin in response to tensile monolayer stress. (A) Effect of Calyculin A on junctional intensity of GFP-Myosin VI ^WT^ or GFP-Myosin VI ^L310G^. (B, C) Effect of Calyculin A on junctional stability (representative FRAP plots) of GFP-Myosin VI ^WT^ (B) or GFP-Myosin VI ^L310G^ (C) (D) Effect of calyculin on total cellular Myosin VI: western blots. (E) Representative E-cadherin/Myosin VI PLA images in control and calyculin-stimulated cells. (F) Western analysis of cell lysates from Myosin VI KD and reconstitution with Myosin VI transgenes (corresponding to Fig 4H). Data are means ± SEM from n=3 independent experiments. ****p<0.0001, ***p<0.001, n.s., not significant; unpaired t-Test (C) with Welch’s correction or 2-way ANOVA (F, H, I) with Sidak’s multiple comparisons test. Scale bars represent 20 μm (B).

**Figure S5 related to Figure 5.**
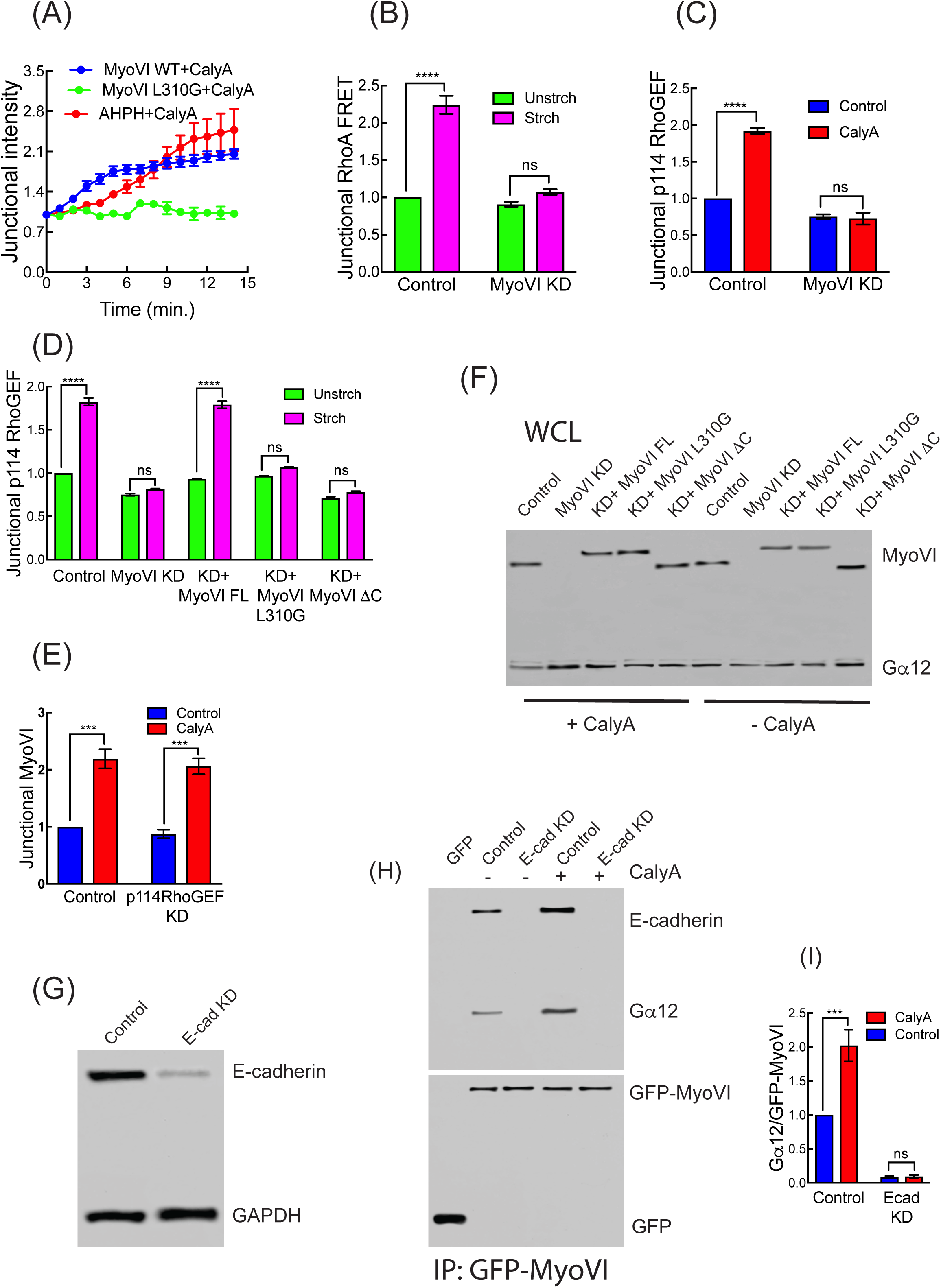
Myosin VI supports tension-activated junctional RhoA signaling. (A) Time course of junctional Myosin VI ^WT^, Myosin VI ^L310G^ and AHPH after stimulation with calyculin (5 movies/conditions/experiment; n=3). (B) Effect of Myosin VI RNAi on junctional RhoA signaling measured by FRET induced by mechanical stretch. (C,D) Effect of Myosin VI RNAi on junctional p114 RhoGEF on stimulation with calyculin A (C) and mechanical stretch (D). (E) Junctional Myosin VI is unaffected by p114RhoGEF KD in calyculin A treated cells. (F) Total cellular expression of Myosin VI and Gα12 on Myosin VI RNAi cells and reconstituted with Myosin VI transgenes; to Fig. 5G. (G) Total cellular expression of E-cadherin after RNAi. (H, I) Effect of E-cadherin RNAi on coimmunoprecipitation of Gα12 with EGFP-Myosin VI: representative blot (H) and quantitation (I). Data are means ± SEM from n=3 independent experiments. ****p<0.0001, ***p<0.001, n.s., not significant; 2-way ANOVA (B-D, I) with Sidak’s multiple comparisons test.

**Figure S6 related to Figure 6.**
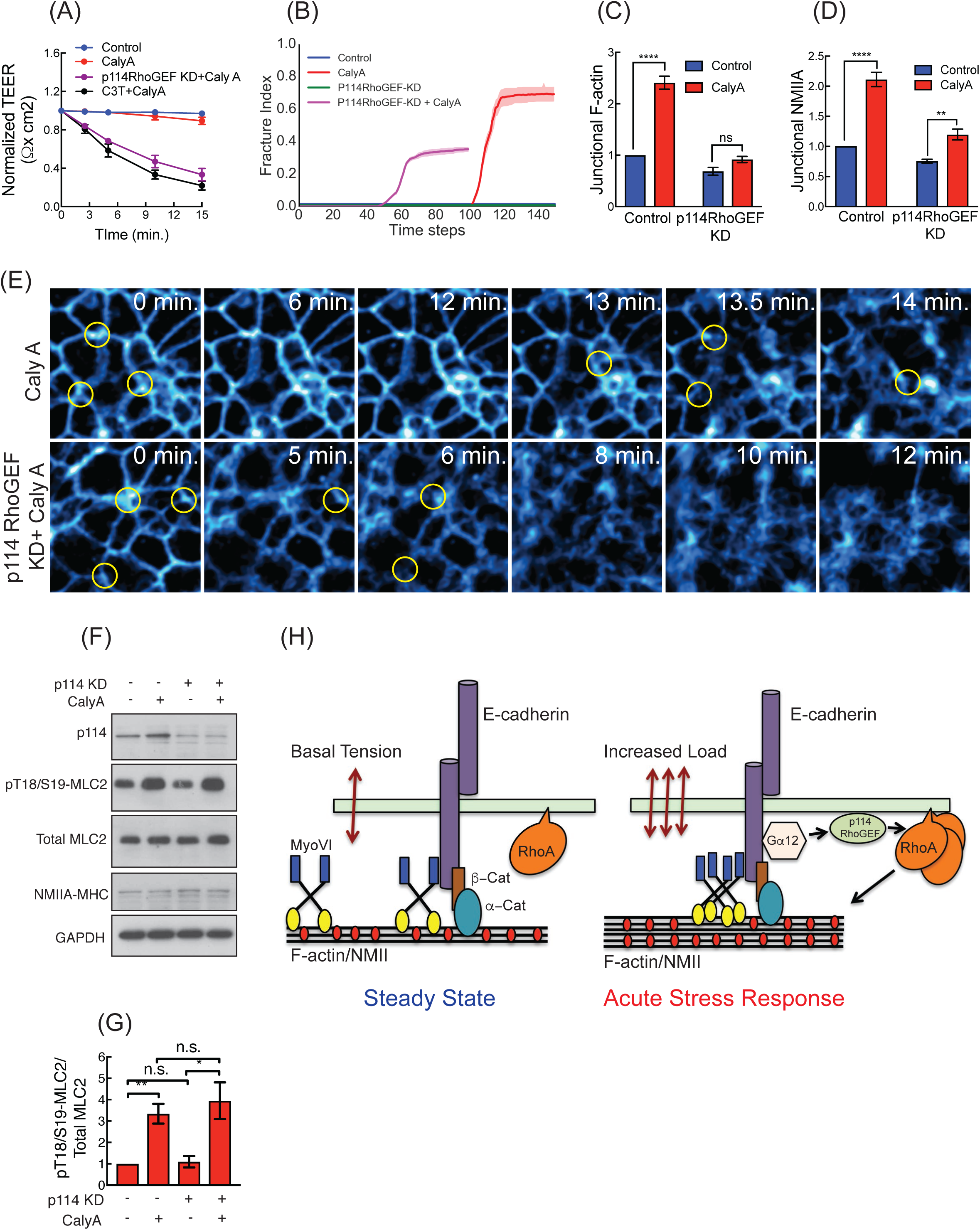
Tension-activated junctional RhoA signaling preserves epithelial integrity. (A)Response of TER to calyculin in p114RhoGEF KD cells or cells treated with C3T. (B)Fracture index for simulated monolayer. Shading shows 95% confidence intervals over three simulations of 800 cells with unique initial seedings. *Δt* = *0.04* t for all simulations, but was recursively reduced if any Δ***F***^*j*^ was greater than 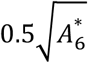 Parameters (*Δϕ*_A_, *Δϕ*_P_, *Y* = 0, 0, 0.92, 0.03, 0.06, 3.46. 0, 0, 0.92, (0.03, 0.03, 0.92) for Control, CalyA, p114 RhoGEF KD and p114 RhoGEF KD+Caly respectively. (C, D) Effect of Calyculin A F-actin (B) and NMIIA (C) at bicellular junctions. (E) Stills from Movie 9 and Movie 10. Yellow asterisks indicating the vertices where fracture occurred. (F,G) Effect of calyculin and p114 RhoGEF KD on cellular levels of active Myosin II (pp-MLC2, pT18/S19) and NMIIA: representative immunoblot (F) and quantitation (G). (H) Cartoon representing the mechanistic model for RhoA activation at AJ in response to acute mechanical stress. Data are means ± SEM from n=3 independent experiments. ****p<0.0001, **p<0.01, *p<0.05, n.s., not significant; One way Anova (D) with Dunnett’s post hoc test or 2-way ANOVA (B, C and F) with Sidak’s multiple comparisons test.

**Figure S7 related to Computational Supplement text.**
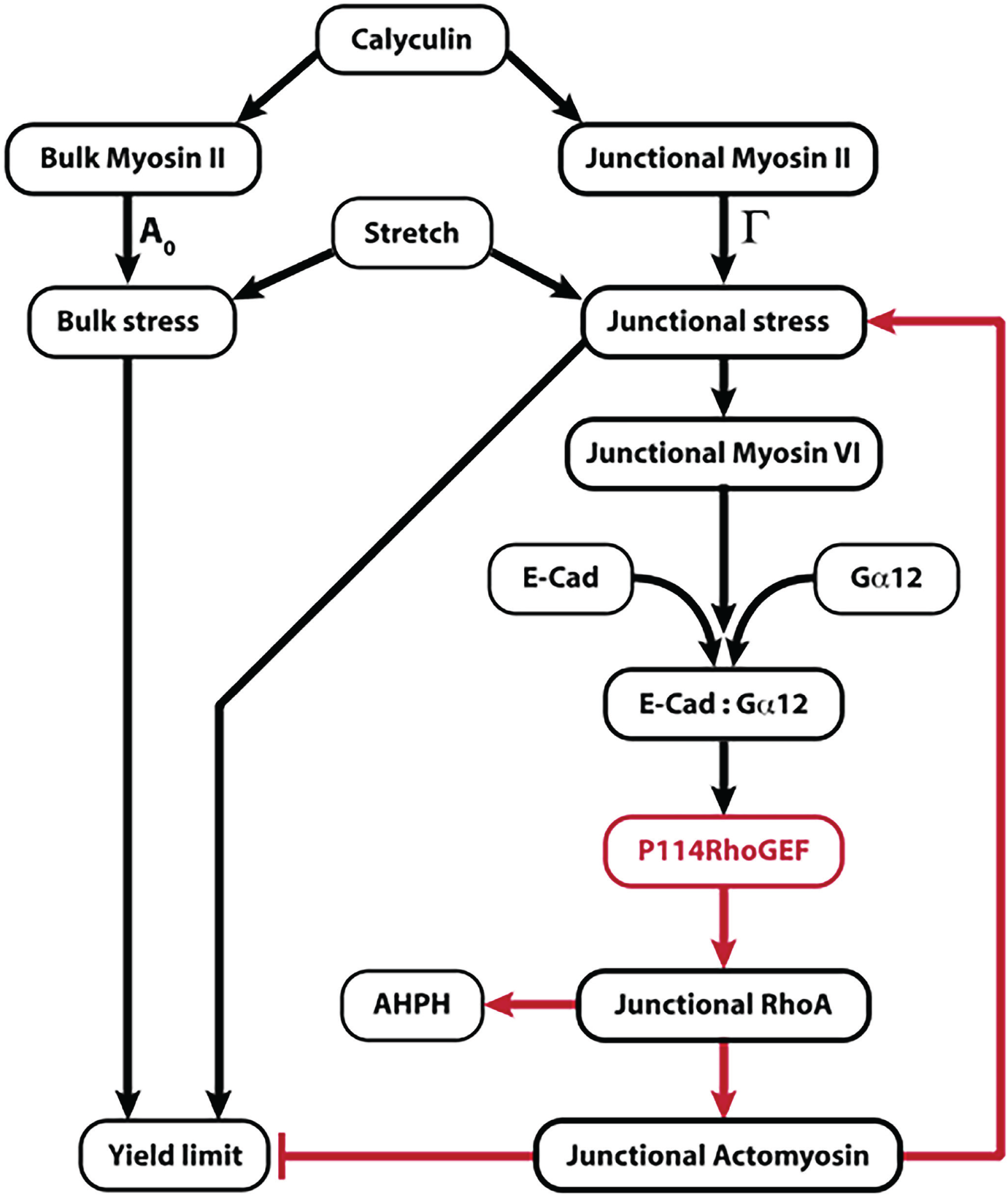
Putative causality network summarising proposed tissue protective mechanism. The p114 RhoGEF mediated RhoA pathway (red) is activated in response to mechanical stress, mechanosensor myosin VI at ZA potentiate the interaction between E-cadherin and Gα12, leading to an increase in the yield limit of cell vertices and an increase in junctional stiffness.

## Supplementary Movie Captions

**Movie S1.** Stress map showing the effect of calyculin A treatment on epithelial monolayer, related to Figure 1. Segmented cells on E-cadherin GFP monolayer showing predicted distribution of effective cell pressures, *P*^eff^. Cells in darker red (blue) are predicted to be under higher net tension (compression). Simulation details are explained in the Computational Supplement.

**Movie S2.** Effect of calyculin A on E-cadherin GFP monolayer, related to Figure 1. Monolayer was imaged for 15 min after adding calyculin (20 nM) and presented at 15 frames/sec.

**Movie S3.** Steady state behaviour of control monolayers, related to Figure 6. Monolayer. Movie was acquired for 15 min after adding DMSOand presented at 15 frames/sec.

**Movie S4.** Steady-state behaviour of p114 RhoGEF RNAi monolayer, related to Figure 6. E-cadherin GFP was imaged for 12 min after adding DMSO, represented at 15 frames/sec.

**Movie S5.** Effect of calyculin A on p114 RhoGEF RNAi monolayer. E-cadherin GFP monolayer knocked down for p114 RhoGEF, related to Figure 6. E-cadherin GFP was imaged for 12 min after adding DMSO, represented at 15 frames/sec.

**Movie S6.** A representative simulation of p114 RhoGEF KD calyculin-treated tissue. The monolayer was randomly initialised with 800 cells satisfying *P*^tis^ = 0 (no net tissue pressure), for parameters (*Λ, Γ*) = (–0.259, 0.186). Calyculin treatment was modelled by decreasing the preferred area and increasing the cortical stiffness at every time step. The yield limit of vertices was not increased in response to stress. Parameters used were (*Δϕ*_A_ = 0.3, *Δϕ*_P_ = 0.3, *Y* = 0.92, *Δt* = *0.04*). Cells in darker red (blue) are predicted to be under higher net tension (compression). Arrows on cell vertices represent magnitude and direction of the net force acting on the vertex.

**Movie S7.** A representative simulation of a tissue that increases junctional stiffness in response to calyculin-induced stress, but does not increase the yield limit of vertices. The monolayer was randomly initialised with 800 cells satisfying *P*^tis^ = 0, for parameters (*Λ,Γ*) = (–0.259, 0.186). Calyculin treatment was modelled by decreasing the preferred area and increasing the cortical stiffness at every time step. Parameters used were (*Δϕ*_A_ = 0.3, *Δϕ*_P_ = 0.6, *Y* = 0.92, *Δt* = *0.04*). Cells in darker red (blue) are predicted to be under higher net tension (compression). Arrows on cell vertices represent magnitude and direction of the net force acting on the vertex.

**Movie S8.** A representative simulation of calyculin-treated control tissue. The monolayer was randomly initialised with 800 cells satisfying *P*^tis^ = 0, for parameters (*Λ, Γ*) = (–0.259, 0.186). Calyculin treatment was modelled by decreasing the preferred area and increasing the cortical stiffness at every time step. The yield limit of multicellular vertices was increased in response to stress. Parameters used were (*Δϕ*_A_ = 0.3, *Δϕ*_P_ = 0.6, *Y* = 3.46, *Δt* = *0.04*). Cells in darker red (blue) are predicted to be under higher net tension (compression). Arrows on cell vertices represent magnitude and direction of the net force acting on the vertex.

**Movie S9.** Fracture pattern of control monolayer treated with calyculin, related to Figure S6. E-cad-GFP cells were imaged for 15 min after adding calyculin; presented at 15 frames/sec.

**Movie S10.** Fracture pattern of p114 RhoGEF KD monolayer treated with calyculin, related to Figure S6. E-cad-GFP cells were imaged for 12 min after adding calyculin; presented at 15 frames/sec.

**Movie S11.** A simulation demonstrating the effect of gradually increasing cortical stiffness, as *ϕ*_+_ increases from 1 to 4 (a fourfold increase in cortical stiffness, *Γ*). The monolayer was randomly initialised with 800 cells satisfying *P*^tis^ = 0, forparameters (*Λ, Γ*) = (–0.259, 0.186). Cells in darker red (blue) are predicted to be under higher net tension (compression). Arrows on cell vertices represent magnitude and direction of the net force acting on the vertex.

## Materials and Methods

### Cell culture and transfection

Human colorectal Caco-2 cells were obtained from ATCC (HTB-37) and cultured in RPMI media supplemented with 10% FBS, 1% non-essential amino acids, 1% l-glutamine and 1% penicillin/streptomycin. Cells were tested for mycoplasma and source cultures maintained in low doses of plasmocin (Invivogen) to prevent mycoplasma contamination. E-Cad-GFP Caco-2 cells generated by CRISPR/Cas9 genome editing were described earlier (Liang, et al., 2017). β-catenin, α-Catenin, and p120-catenin immunoprecipited with E-cadherin GFP in the working clone (B1C2) to the similar extent as did E-Cadherin from a control Caco-2 line cells (Liang, et al., 2017). Quantitative immunofluorescence also showed identical levels of E-cadherin, α-Catenin, NMIIA and F-actin in the E-Cad-GFP Caco-2 line as in control Caco-2 cells. Cells were transfected at 50-60% confluency using Lipofectamine 3000 (Invitrogen) for expression constructs or RNAiMAX (Invitrogen) for RNAi oligonucleotides according to the manufacturer’s instructions. Cells were processed for experiment and analysis 24-48 h post transfection as required.

### Application of equibiaxial static stretch

Caco-2 monolayers was grown on collagen-coated 25 mm BioFlex culture plates. Cells were subjected to static stretch using a Flexcell Fx-5000TM Tension System (Flexcell International, Hillsborough, NC) for 10 min with 10% strain. Control wells were plugged at the bottom by rubber capping without application of any stretch. The inhibitors were present during the static stretch wherever needed.

### siRNA, Antibodies and Drugs

Target proteins were depleted by RNAi in Caco 2 cells using custom designed or commercially available siRNAs (Invitrogen, Sigma USA or IDT, Singapore). Relevant siRNA sequences are shown in Supplementary Table 2. Algorithms from Dharmacon were used to generate RNAi sequences for custom designed siRNA targeting the ORF or the 3’UTR.

Primary antibodies used in this study were: rabbit pAb against Phospho-myosin light chain 2 (Cell Signaling: Cat#3674, 1:200 IF, 1:1000 WB), mouse mAb against Myosin light chain 2 (Abcam: Cat#ab89594, 1:1000 WB), rabbit pAb against α-Catenin (Invitrogen #71-1200, 1:100 IF, 1:1000 WB); rat pAb α-18 (a kind gift from Akira Nagafuchi, Nara Medical University, Japan, 1:200 IF), goat pAb against p114RhoGEF (Abcam: Cat#ab10152, 1:50 IF, 1:250 WB), rabbit pAb against p114RhoGEF (Abcam: Cat#ab96520, 1:100 IF, 1:1000 WB), rabbit pAb against Gα12 (Abcam: Cat#ab154004, 1:100 IF, 1:500 WB), rabbit mAb against Gα13 (Abcam: Cat#ab128900, 1:100 IF, 1:500 WB), rabbit pAb against GEFH1 (Abcam: Cat#ab155785, 1:100 IF, 1:1000 WB), mouse mAb against LARG1 (Merck: Cat#MABT124, 1:100 IF, 1:1000 WB), rabbit pAb against ECT2 (Merck-Millipore: Cat#07-1364, 1:50 for IF, 1:500 WB), mouse mAb against E-cadherin (clone HECD-1, a kind gift from Peggy Wheelock, University of Nebraska, USA, with permission from Masatoshi Takeichi, 1:1000 WB); rat mAb against E-cadherin (Invitrogen: Cat#191900, 1:500 IF), mouse mAb against Myosin IIA (Abcam: Cat#ab55456, 1:250 IF); rabbit pAb against Myosin IIA (WB only, Sigma: Cat#M7939, 1:2500); rabbit pAb against Myosin IIA (Covance: Cat#PRB-440P, 1:100 IF); mouse mAb against Myosin IIB (Abcam: Cat#ab684); rabbit pAb against Myosin IIB (WB only, Sigma:Cat#M8064, 1:2500), Myosin IIB (Covance: Cat#PRB-445P, 1:100 IF), rabbit pAb against Myosin VI (for IF only 1:200, (Maddugoda, et al., 2007)), mouse mAb against Myosin VI (WB only, Sigma: Cat#M0691, 1:250), mouse mAb against GFP (Roche: Cat#A11120, 1:500 IF, 1:5000 WB), Phalloidin conjugated with AlexaFluor546 or AlexaFluor647 (Invitrogen; 1:250) was used to label F-actin.

The species-specific secondary antibodies used for immunofluorescence in this study were conjugated with AlexaFluor 488, 594 and 647 (Invitrogen, 1:500) or with horseradish peroxidase conjugated secondary antibodies (Bio-Rad Laboratories, 1: 5,000/10,000) for immunoblotting.

Cells were treated with Calyculin A (ab141784, Abcam, USA; 20 nM, 12-15 min) or Y-27632 (no. Y0503, Sigma; 30 μM, 1 h). For RhoA activity inhibition, cells were treated with 1 μg/ μl of cell-permeable RhoA inhibitor (C3-T based; no. CT04-A, Jomar Bioscience) for 1h before Calyculin A addition.

### Plasmids

The GFP–AHPH location RhoA biosensor was a kind gift from M. Glotzer (University of Chicago, USA. The CFP-p114RhoGEF construct is a kind gift from K. Mizuno, Tohoku University; Japan. This construct was used as a template for generating p114RhoGEF-Y260A mutant by using the Quick Change V Site Directed Mutagenesis Kit (New England Biolabs, USA) as per the manufacturer’s protocol and the corresponding primers are listed in Supplementary Table 5. Untagged version of G protein-alpha 12 (Q231L; #46825) and pTriEx-RhoA FRET WT biosensor (#12150) were obtained from Addgene. Untagged G protein-alpha 12 (Q231L) construct was used as template to PCR amplify the full-length protein and was subsequently cloned into pEGFP-C1 (Clontech) using EcoR1 and BamH1 restriction sites. For Gα12ΔEcad construct, we PCR amplified fragment-1 (1-130 aa) and fragment 2 (197-365 aa) and were subsequently cloned in to pEGFP-C1 (Clontech) using EcoR1/Knp1(fragment 1) and Knp1/BamH1 restriction sites. Porcine GFP-myosin VI construct was a gift from Dr. T. Hasson (University of California, San Diego, USA). This construct was used as the templatesfor generating GFP-myosin VI L310G mutant with the the Quick Change V Site Directed Mutagenesis Kit (New England Biolabs, USA) as per the manufacturer’s protocol; relevant primers are listed in Supplementary Table 2. For GFP myosin VI-ΔC construct, a PCR amplified fragment (1-1020 aa) from full-length GFP myosin VI was subsequently cloned in to pEGFP-C1 (Clontech) using EcoR1 and BamH1 restriction sites. For αE-Cat TS, the mouse N-terminal (1-697 aa) of αE-Catenin was inserted before mTFP1 by in-fusion cloning (Takara Bioscience, Clonetech) at the Xho1 restriction site. The C-terminal fragment (698-906 aa) of αE-Catenin was similarly cloned after VenusA206K at the Not1 restriction site. The details of construction and validation of the αCat-TS construct are described elsewhere (Acharya, et al., 2017).

### Immunofluorescence and live-cell microscopy

For immunofluorescence, cells were either fixed with methanol at −20°C or with 4% paraformaldehyde in cytoskeleton stabilization buffer (10 mM PIPES at pH 6.8, 100 mM KCl, 300 mM sucrose, 2 mM EGTA and 2 mM MgCl_2_) and subsequently permeabilized with 0.25% Triton-X in PBS. Upright Zeiss LSM 710 Meta scanning microscopes (63X, 1.4NA Plan Apo objective) driven by Zen software (ZEN 2012, Zeiss) were used for fixed imaging. Zeiss ELYRA PS.1 SIM/STORM microscope (63X, 1.4NA Plan Apo objective) with sCMOS camera driven by Zen software (ZEN 2012, Zeiss) was used for SIM imaging. All image reconstruction and channel alignment were performed within the ZEN software.

For live cell imaging, FRAP and FRET experiments, the cells were grown in 29 mm diameter borosilicate glass-bottomed dishes (Shengyou Biotechnology) and imaged in movie medium (15 mM HEPES, pH 7.5; 5 mM CaCl2; 10 mM D-glucose; 2.5% FBS in Hank’s balance salt solution) at 37 °C. GFP-E cadherin movies and FRAP was conducted on Inverted Zeiss LSM 710 Metal NLO AiryScan confocal system. For FRAP, either 488 laser or Mai-Tai-eHP multiphoton laser (2000mW laser power) with dedicated BiG(GaAsP) detectors used. FRET analysis was conducted using LSM 710 Meta confocal with BiG(GaAsP) detector for CFP/YFP imaging.

### Image processing and analysis

Quantitative analysis of junctional intensity of E-cadherin, NMIIA, NMIIB, F-actin, αE-Catenin and α18 was performed in ImageJ software (NIH) with the line scan functions by drawing a line of 12 μm in length (10-pixel wide) orthogonal to, and centred on, randomly selected homotypic junctions. Optical Z-stacks (0.2 μm intervals) were acquired to correct for cell heights and to focus on all junctions analyzed. The pixel intensities along the selected line was recorded and plotted, the fluorescence profiles were fitted to a Gaussian curve and the peak values (fluorescence intensities at junctions) were obtained from this fitting. The average pixel intensity values lying on either side of the center (at junctions) on the masking line were considered as background fluorescence and subtracted from the plots. To correct for fluctuations in the height of cells within frames, some representative confocal images (identified in figure captions) are presented as maximum projection views of the three most-apical sections of the cells (0.19 μm intervals between the sections). The junctional intensity of α18 is presented as the ratio of junctional α18 and junctional αE-Catenin. To quantify GFP-AHPH, Gα12 and different RhoA GEFs we measured the ratio of junctional intensity and cytoplasmic mean fluorescence intensity in ImageJ. A 10-pixel wide line (using the freehand drawing tool) was drawn covering the region of whole junctions, and the mean pixel intensity of this region was measured as the junctional GFP-AHPH. For GEFs and Gα12, the masking was done on the corresponding E-cadherin in the other channel of the same image. Mean cytoplasmic GFP-AHPH, GEFs and Gα12 fluorescence was measured by masking the entire cytoplasm of cells and calculating the average pixel intensities within that mask. Data are presented as the ratio values normalized to the corresponding ratio value for control conditions and the normalized ratio of junctional versus cytoplasmic fluorescence intensity is referred to as Junctional AHPH, Junctional GEFs or Junctional Gα12.

For PLA analysis, we have masked the junction using the F-actin staining and measured the PLA dots on that masked region for all images using ComDet plugin of ImageJ. To analyse the fluorescence intensity at vertices, a constant circular ROI of 3.1 μm diameter was drawn encircling the vertices. The mean intensity of all pixels within that ROI was calculated and plotted for E-cadherin and F-actin in different experimental conditions.

To quantitate fracture initiation from movies, we measured the change in number of intact vertices (as fractures were overwhelmingly initiated at vertices). We measured the number of intact vertices present at each sample time point (using Tissue Analyzer and ComDet plugin of ImageJ), subtracted this from the total number of vertices present at the start of the movie, and normalized this difference value to the initial number of vertices. For junctional fracture index, the entire image was segmented into small square templates and the number of junctions were counted.

### FRET and FRAP Analysis

For αE-Cat TS FRET (Acharya, et al., 2017), human αE-Catenin specific siRNA transfected Caco-2 cells were reconstituted with αE-Cat TS by transient transfection and processed for FRET experiments after 24 h. Cells were imaged live (Calyculin A) at 37°C of fixed (Stretched membrane) by confocal microscopy. Images were acquired by sequential line acquisition. Donor (mTFP1 in CFP channel) and FRET channels were recorded by scanning using a 458nm laser; the emission was collected in the donor emission region (BP 460–490 nm) and acceptor emission region (BP 520–560 nm), respectively. The acceptor (VenusA206K in YFP channel) was imaged using a 514 nm laser line for excitation and emission was collected in the acceptor emission range (BP 530–560 nm). FRET index was calculated pixel by pixel as the ratio between the FRET channels and acceptor channel. Data showing the normalized FRET index compare to untreated control. For RhoA-FRET, Caco-2 cells were transfected with control or respective siRNA 24 h before the transfection of pTriEx-RhoA biosensor. FRET measurements and analysis were performed 36 h after biosensor transfection following mechanical stretch (fixed cell) or calyculin A treatment (live at 37°C). The Donor (CFP) and FRET channels were excited using the 458 nm laser line and emissions were recorded between 470 and 490 nm (Donor) and between 530 and 590 nm (Acceptor, YFP), respectively. The Acceptor channel was excited using the 514 nm laser line and the emission was recorded between 530 and 590 nm. The average FRET/YFP emission ratios were calculated as FRET index on a pixel-by-pixel basis at the apical junctions. The linear pixels within the ROIs of both YFP and CFP channels at apical junctions were determined and only included them in FRET calculation as described earlier (Ratheesh, et al., 2012). Mean FRET/YFP ratio of junctional pixels of three independent experiments were normalized with untreated or unstretched control and plotted for statistical analysis. We determined the linear pixels in our images by calculating the emission ratio of FRET/YFP and FRET/CFP for every pixel present within the ROIs across all of the images of the control condition. We therefore sorted all pixels according to Acceptor and Donor intensity values. For each intensity value, an average FRET emission ratio was calculated using a custom-made MATLAB script. We plotted the average FRET emission ratios for FRET/YFP and FRET/CFP against Donor and Acceptor intensities, respectively. Non-linear behaviour was seen as an overflow of the FRET emission ratio (FRET/YFP or FRET/CFP), and an appropriate threshold of intensity values was determined to exclude the pixels within this non-linear region before further analysis.

For FRAP analysis. Caco-2 cells were transfected with siRNA targeted against either the UTR region of p114RhoGEF, Gα12 or ORF of Myosin VI. 24 h after the respective KD cells were transfected with GFP-p114RhoGEF, GFP-Gα12 and GFP-Myosin VI WT or L310G for FRAP experiment (24 h post transfection). For E-cadherin FRAP, both junctional and vertices we used E-cadherin GFP stable line. To obtain FRAP profiles, a constant circular ROI was drawn in the centre of the junction (or encircle the vertices) and was bleached to 70–80% with 810 nm laser (for p114RhoGEF, Gα12 and Myosin VI; at 26%, 1 iteration). For E-cadherin, 488 nm laser at 100% transmission was used for photobleaching. Time-lapse images were acquired before (pre-bleach) and after bleaching with a constant interval. The average fluorescence intensity F(Tt) over time was in bleached area was analysed with ImageJ. The mean values of all frames before bleaching was used as the pre-bleached value F(Tp). The value of the first frame after bleaching was defined as F(To). FRAP values were the calculated as:

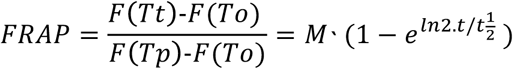

the calculated recovery fluorescence was plotted over time, where Mf is the mobile fraction, t1/2 is the half time of recovery and Tt is time in seconds. The FRAP values were fitted using a nonlinear regression and the exponential one or two-phase association model (only for junctional p114RhoGEF) using Y0 = 0 and where Mf corresponds to the plateau value in Prism software. The immobile fraction was then calculated as:

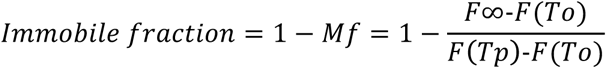

10 junctions were measured in each experiment (n=3). Mean ratios of independent experiments were plotted and used for statistical analysis.

### Nematic Order analysis of F-actin

F-actin filament orientation was assessed with the nematic order parameter (Reymann, et al., 2016) using the technique and Matlab codes of Reymann et al (2016) based on Fourier Transform Analysis of phalloidin staining imaged by SIM. The entire junction was randomly segmented in several square templates of 2.64 micrometer (N ∼ 25) for quantitation. For vertices, one square template covers the entire vertex.

### Immunoprecipitation and Immunoblotting

For Gα12 and Myosin VI GFP-trap, Caco-2 cells were cultured on 10 cm culture dishes, at 50-60%% confluence for transfection with siRNA (wherever necessary) using lipofectamine RNAiMax and 24 h after the GFP tagged expression constructs (∼20 μg) was transfected with Lipofectamin 3000 (Invitrogen). The clarified cell lysate was incubated with GFP-trap beads. Following incubation, GFP-trap beads were washed several times in lysis buffer supplemented with 300 mM NaCl. The protein complexes were then resolved by SDS-PAGE and immunoblotted. For E-cadherin endogenous immunoprecipitation, 1mg of total protein was added to 200 μl of culture supernatant of E-cadherin monoclonal antibody. Following overnight rotation, the lysate-antibody solution was incubated packed slurry of Protein A. Protein A Beads were then washed three times on ice to remove non-specific binding, boiled in the SDS-PAGE loading buffer and then centrifuged at 12,500 g; 10 min. The supernatants were subjected to Western blotting.

### Transepithelial electrical resistance (TEER) measurements

Caco 2 cells were plated at a 2×10^5^ on Transwells with polyester membrane inserts (12 mm diameter, 0.4 mm-pore size; Corning, MA). TEER was measured using a Millicell-ERS epithelial volt-ohmmeter (Millipore, Billerica, MA). TEER values (ohms.cm2) were normalized based on the area of the monolayer and was calculated by subtracting the blank values from the filter and the bathing medium. For C3T treatment, the cells were pre-incubated for 45 min. with C3T and then DMSO or calyculin was added for 15 minutes and TEER was measured for indicated time points after 21 days culture the cells. For p114 RhoGEF KD, the cells were trypsinized and resuspended with Opti-MEM. Lipofectamin RNAiMAX with control or p114 RhoGEF specific siRNA was mixed and incubated for 20 min. Then the transfection mix was mixed with cell suspension and overlayed on apical chamber of the insert. The bottom chamber was filled with Opti-MEM as well. 24 h after, the media was replaced with fresh RPMI media without antibiotic. Once cell seeded, in every 3 days, the similar siRNA transfection mix was made with Opti-MEM and RNAiMAX added to apical chamber of insert and as well as in bottom well and incubated for 6 h, followed by media replacement with RPMI without antibiotic. The last transfection was done on day 18^th^ and the TEER was measure on 21^st^ Day. On the day of experiment, the DMSO or calyculin was added for 15 minutes and TEER was measured for indicated time points.

### Statistical Analysis and Code Availability

All data displayed are represented as mean ± SEM, derived from three or more independent experiments as indicated in figure legends, and all statistical analyses were performed using GraphPad Prism. Unpaired two-tailed t tests were used to compare datasets consisting of two groups, and a Welch’s correction was included when data normalized to the control values were being assessed. One-way ANOVA with Dunnett’s post hoc test was used to compare three or more groups. For comparing two different independent conditions with three or more groups two-way ANOVA with with Sidak’s multiple comparisons test was performed. The MATLAB scripts for FRET analysis as the codes for simulations of the vertex model described in this paper are available on request.

